# ChemChaste: Simulating spatially inhomogenous biochemical reaction-diffusion systems for modelling cell-environment feedbacks

**DOI:** 10.1101/2021.10.21.465304

**Authors:** Connah G. M. Johnson, Alexander G. Fletcher, Orkun S. Soyer

## Abstract

**Motivation:** Spatial organisation plays an important role in the function of many biological systems, from cell fate specification in animal development to multi-step metabolic conversions in microbial communities. The study of such systems benefits from the use of spatially explicit computational models that combine a discrete description of cells with a continuum description of one or more chemicals diffusing within a surrounding bulk medium. These models allow the *in silico* testing and refinement of mechanistic hypotheses. However, most existing models of this type do not account for concurrent bulk and intracellular biochemical reactions and their possible coupling.

**Results:** Here, we describe ChemChaste, an extension for the open-source C++ computational biology library Chaste. ChemChaste enables the spatial simulation of both multicellular and bulk biochemistry by expanding on Chaste’s existing capabilities. In particular, ChemChaste enables: (i) simulation of an arbitrary number of spatially diffusing chemicals; (ii) spatially heterogeneous chemical diffusion coefficients; and (iii) inclusion of both bulk and intracellular biochemical reactions and their coupling. ChemChaste also introduces a file-based interface that allows users to define the parameters relating to these functional features without the need to interact directly with Chaste’s core C++ code. We describe ChemChaste and demonstrate its functionality using a selection of chemical and biochemical exemplars, with a focus on demonstrating increased ability in modelling bulk chemical reactions and their coupling with intracellular reactions.

**Availability and implementation:** ChemChaste is a free, open-source C++ library, available via GitHub at https://github.com/OSS-Lab/ChemChaste under the BSD license.

## 1 Introduction

Understanding the emergent dynamics of spatially heterogeneous cell populations is highly relevant to both eukaryotic and microbial biology. Spatially self-organised biological systems often display nonlinear dynamics (An *et al*., 2017; Hart *et al*., 2019; Painter, 2019), which may be difficult to mechanistically explain through observation alone, necessitating the use of computational modelling approaches to help guide and explain experimental studies. Several outstanding challenges must be addressed to fully leverage models of spatially organised biological systems (Fletcher and Osborne, 2021), not least the development of robust and extensive computational frameworks that allow users to define, explore, and share models in a straightforward manner.

Many computational frameworks already exist for studying the dynamics of spatially organised cell populations. Some of these, such as iDynoMiCs (Lardon *et al*., 2011), use a bottom-up (discrete, agent-based) approach to modelling individual cell behaviours (Kreft *et al*., 2017), combined with a top-down (continuum, partial differential equation (PDE) based) approach to modelling the diffusive transport of nutrients and other chemicals. In this approach, some aspects of cell physiology are ‘hard-coded’, along with specific ‘rules’ governing their dynamics. In other computational frameworks, the physical forces acting on individual cells are modelled explicitly, but cell physiology is not. In these approaches, cells are treated as extended shapes in space, with cell proliferation and migration implemented through neighbourhood update rules, e.g. an implementation of the so-called cellular Potts model (e.g. as done in CompuCell3D (Glazier and Graner, 1993) and as used in Morpheus (Starruß *et al*., 2014)). It is also possible to combine these two approaches, into what we call a ‘hybrid continuum-discrete approach’, where cells are represented by particles, with some aspects of their physiology encoded by rules (e.g. cell division) and others governed by spatially explicit energy or force equations (e.g. cell migration). Such hybrid approaches have been developed by either creating dedicated, new computational frameworks (e.g. HAL (Bravo *et al*., 2020), PhysiCell (Ghaffarizadeh *et al*., 2018), Chaste (Cooper *et al*., 2020)), or by adapting existing agent-based (Xavier *et al*., 2005) or molecular dynamics (Plimpton, 1995) tools.

Using hybrid modelling tools, cell physiology can theoretically be coupled to the dynamics of chemicals in the bulk medium. This functionality, however, is implemented in a limited fashion in existing platforms. For example, in Chaste, PhysiCell and CompuCell3D, either only a limited number of bulk chemicals can be dynamically modelled, and/or diffusion coefficients are assumed to be homogeneous. Additionally, the linking of these bulk chemicals to intracellular reactions is limited in terms of number of reactions and couplings that can be encoded in each cell and at the cell-bulk interface. This limits the range of biological phenomena that can be studied within existing computational frameworks.

The coupling between cells and their microenvironment is increasingly being recognised as playing a fundamental role in cell dynamics in the context of both microbial and eukaryotic populations, e.g. metabolic environmental feedbacks in the tumour microenvironment (Carmona-Fontaine *et al*., 2017) and microbial community stability (Ratzke and Gore, 2018). Additional feedbacks can emerge from cell-excreted enzymes, which introduce reactions in the bulk, and from cell-excreted metabolites or proteins that can affect chemical diffusion coefficients in the bulk or near cells. Such effects arising from bulk-cell interaction can create their own nonlinear dynamics (Kondo and Miura, 2010; Newman, 2016; Höfer *et al*., 1995; Glock *et al*., 2019) or exert a feedback onto cellular physiology (Liu *et al*., 2015; Bocci *et al*., 2018; Mikami *et al*., 1992). Thus, modelling of metabolic and other feedbacks between bulk environment and cellular behaviours would benefit from the further development of computational frameworks centred on the role chemical coupling.

To this end, we introduce ChemChaste, a computational framework that allows the simulation of any number of chemical reaction-diffusion systems with or without cells, and allows cell-excreted chemicals or enzymes to react in the bulk phase. ChemChaste builds upon Chaste and expands its capabilities with the introduction of: (i) unlimited number of PDEs for modelling any number of bulk chemicals diffusion dynamics,; (ii) heterogeneous diffusion rates, allowing for implementation of different ‘domains’ in the bulk pertaining different diffusion properties; (iii) expansion of cellular network reaction size that can be implemented to describe cellular behaviours; and (iv) a user-interface for defining model structure. The user-interface allows cell-internal biochemical reaction systems (cell network ODEs), spatial reactions in the bulk, and heterogeneous diffusion rates for chemicals in the bulk to be encoded in a file-based system. These features allow easier simulations in ChemChaste, without any need for users to change the C++ source code. Below, we demonstrate the ChemChaste implementation and functionality using a set of chemical and biochemical exemplars, including a cell-based example. All of the source code and user manuals for ChemChaste are provided through GitHub (https://github.com/OSS-Lab/ChemChaste) as an open-source library to accompany Chaste, allowing for its application and further development by the research community.

## 2 Methods

ChemChaste builds from Chaste, inheriting its adaptable and modular C++ structure (Mirams *et al*., 2013; Cooper *et al*., 2020), and expanding its capabilities with a comprehensive set of C++ classes (Figure 1). Chaste exhibits many capabilities ideal for the foundation of a hybrid modelling framework, including: (i) implementation of a range of on-lattice and off-lattice multicellular modelling approaches in a consistent computational framework (Osborne *et al*., 2017); (ii) centre-based cell modelling, which treats cells as point particles with radii of interactions (Pathmanathan *et al*., 2009); (iii) accounting for cell physiology through empirical rules or a limited intracellular reaction network implemented as a set of ordinary differential equations (ODEs); (iv) modelling of cell physics, including movement and attachment;and (v) modelling of bulk chemicals dynamics using PDEs solved numerically using the finite element (FE) method (Osborne *et al*., 2017). For specific biological modelling applications, Chaste requires the PDEs and ODEs to be explicitly written by the user as C++ classes, limiting Chaste’s usability to those familiar with C++ (Fletcher *et al*., 2013; Dunn *et al*., 2013; Figueredo *et al*., 2013)

**Figure 1:**
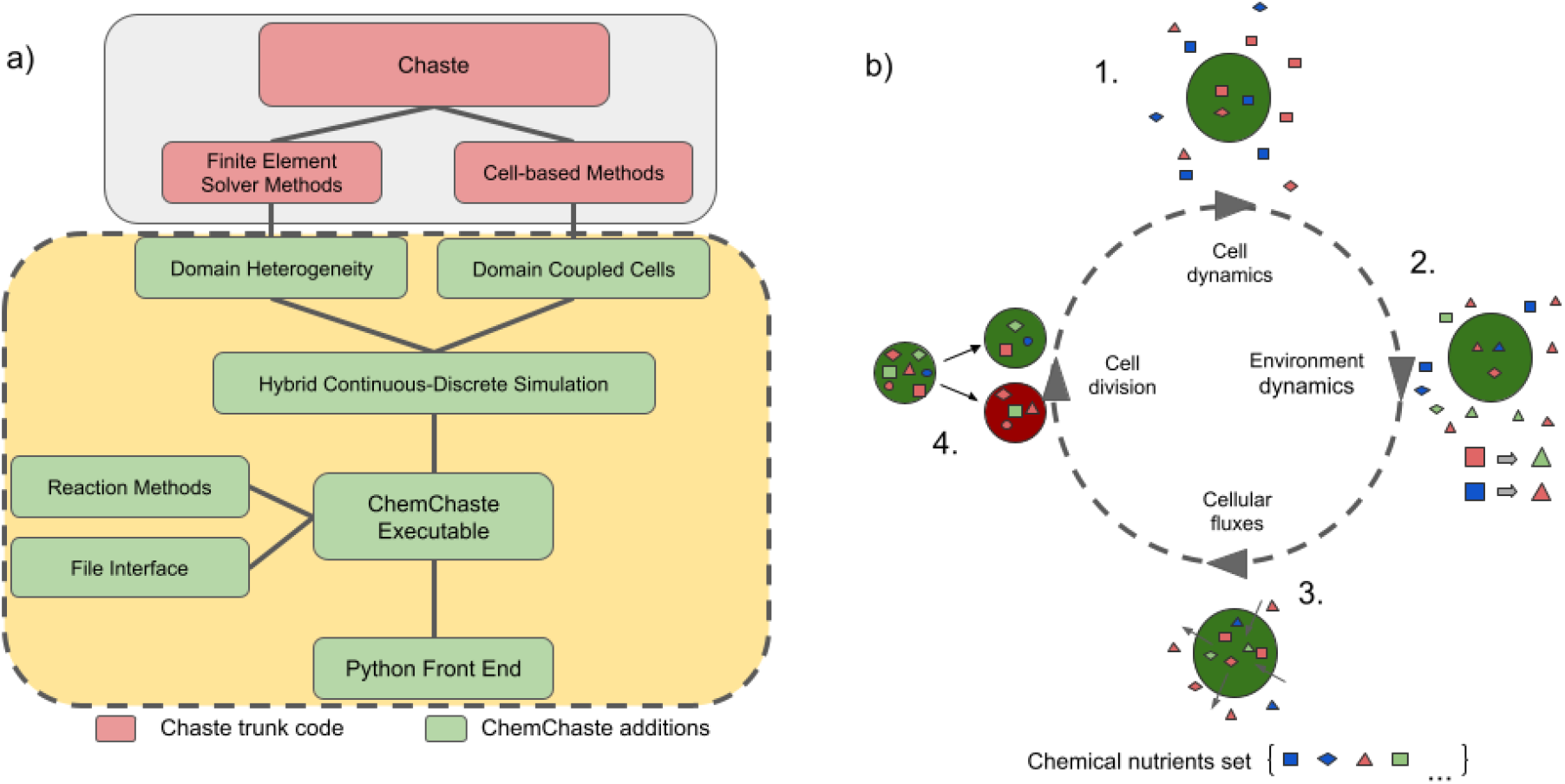
ChemChaste’s simulation framework. a) ChemChaste classes (in dashed yellow-green) that extend Chaste’s FE solver capabilities. These build on existing Chaste modules (in solid gray-pink) and allow for heterogeneous spatial domains with varying diffusion rates for chemicals. The cell-based methods are also extended through introducing transport properties linking cell interior and exterior state variables. These extensions are coupled with a file-based user interface allowing higher-level model specification. b) Four processes that occur over discrete time steps and allow the simulation of cells coupled to the bulk. The cells perform their own system of rules or reactions (cell cycle progression, cell properties, and cellular reaction networks) (1 & 2) before the environmental reaction-diffusion systems are solved (2). The state variables are then coupled through cellular flux (3), before any implemented cell-based rules (e.g. relating to cell division and/or death) are performed (4).

Expanding from Chaste, ChemChaste considers parabolic reaction-diffusion systems, where chemicals diffusing and reacting in the bulk are also coupled with cells present in the same bulk, through cellular excretion and uptake. For simulating such cell-bulk coupling, ChemChaste is developed to handle different chemical species confined to the bulk, to cell populations, or present in both phases. ChemChaste also allows for spatially varying chemical diffusion coefficients.

Each ChemChaste simulation features four distinct dynamical components that run at each discrete time step of the simulation (Figure 1-b). These involve updating of bulk and cellular chemical systems, their couplings, cell behaviours, and cell positions. The bulk and cellular chemical reaction systems are considered separately: the former is updated by solving reaction-diffusion equations, taking into account any reactions implemented in the bulk; while the latter may in general differ from the bulk chemical system and may involve further chemical species. These two systems are coupled through transport of chemicals across the cell membrane. Thus, bulk chemical concentrations are updated according to these couplings. After all chemical concentrations have been updated, any ‘rules’ implemented regarding cell behaviour (e.g. division) are checked and subsequent cellular events (e.g. cell death, division) are implemented. Division introduces a daughter cell into the simulation. In this case, the cellular chemicals of the parent cell are re-distributed between both cells, based on a user-defined parameter (allowing for symmetric or asymmetric inheritance of cellular chemicals). The location of each cell is updated by numerically integrating its equation of motion. These two steps, division and movement, are inherited from Chaste (Cooper *et al*., 2020). The user may tailor the simulation details through a file interface system. Further details of the ChemChaste platform are explained below and in the Supplementary Information (SI).

### 2.1 Expanding the reaction-diffusion system simulations: The Domain Field Class

The core of Chaste is composed of finite element (FE) solvers and associated spatial meshing routines (see SI section S1 for details of the FE method as implemented in Chaste). In brief, the FE methods model the bulk domain as a discrete mesh of nodes and approximates the concentration of each chemical across this mesh, subject to a user-defined combination of boundary conditions (BCs): Neumann; Dirichlet; or periodic conditions at the edge of the bulk domain. Over the mesh, Chaste utilises a range of ODE solvers, chosen by the user, to determine the ODE solutions at the discrete mesh nodes. Utilising a set of linear basis functions, these nodal ODE solutions are then interpolated onto a finer grid of points, known as Gauss points, where point-based source terms and diffusive terms are added. Chaste’s FE method then uses the chemical values at the Gauss points to compute the PDE system solutions at the next time step. This implementation has been limited in Chaste to solving the same given ODE for all nodes in the mesh.

Expanding from this implementation, ChemChaste introduces a `Domain Field` class, which allows us to compute the solution of nodal ODEs generally varying at each mesh node. With the addition of the chemical and reaction classes (see SI, sections S1.3–S1.4), ChemChaste forms a chemical `Domain Field` wherein the concrete reaction systems are mapped to the FE mesh. This expansion allows for: (i) multiple, diffusing bulk chemicals; (ii) reactions among chemicals in the bulk; and (iii) spatially varying diffusion rates for chemicals. With this introduction the simulation domain may now be broken into sub-domains, each containing their own diffusion parameters, ODE systems, and node-based source terms. This allows chemical reaction systems to be confined to sub-domains of the simulation for modelling spatial sub-compartments with their own diffusion parameters, e.g. a biofilm or tissue surrounded by a bulk. The `Domain Field` class uses a 2D matrix to contain the nodal values which acts as a look up reference for spatial aspects of the simulation. While this currently limits the ChemChaste simulation to a 2D domain, an extension to 3D simulations would be straightforward for a C++ proficient user by editing the source code.

### 2.2 Coupling the cell physiology and reaction-diffusion system simulations

The core spatial mesh routines of Chaste also form the basis of simulating dynamic cell populations. ChemChaste uses the ‘node-based’ or cell-centre modelling approach offered in Chaste (Pathmanathan *et al*., 2009). In this approach, a cellular mesh (CM) is defined wherein each mesh node acts as the centre of a cell. Each cell is simulated as a particle, and the CM vertices are used to encode any rules (e.g. physical forces) governing physical cell interactions (Osborne *et al*., 2017; Fletcher *et al*., 2013). The CM is also mutable, allowing simulation of cell motility - by defining forces to shift CM nodes - or cell division and death - by performing vertex additions or deletions on the CM (Mirams *et al*., 2013).

ChemChaste expands upon this node-based cell population simulation to introduce the coupling between cellular and bulk chemicals. As explained above, an interpolated Gauss point is produced during the FE simulations. In ChemChaste, this point may also be the location of a cell in the CM. When this is the case, membrane and transport reactions are performed on the selected cell and their outcomes are coupled to the relevant cellular and bulk chemicals. In this way the cell’s ‘contribution’ to the source term of the related, bulk chemicals’ reaction-diffusion PDE is accounted for. At the same time the selected cell’s internal chemical concentrations are updated through exchanged chemicals (see SI, section S1.2).

### 2.3 Specifying chemical reactions and chemicals diffusion properties

ChemChaste allows modelling of three different reaction processes based on where they occur; bulk, membrane, and transport reaction. Bulk reactions offer the means to model reactions in the bulk and acting on spatially diffusing chemical species. As explained above, the FE simulations implement on each node of the mesh a reaction rule, which is used to update species’ concentrations accordingly. Bulk reactions occur on these mesh nodes and act as a source/sink term for the PDEs defining the reaction-diffusion system. Membrane and transport reactions involve cellular and bulk chemical species and therefore require knowledge of the concentrations of a given chemical both within the cell object and in the bulk. In the case of membrane reactions, reaction rates depend on both bulk and intracellular chemical concentrations, however, there is no chemical species exchange through the membrane. This class of reactions is thus ideal for implementing processes such as membrane bound enzymatic reactions. Transport reactions implement a chemical flux through the membrane and internal species may react or exchange with external species.

The three reaction types are modelled with user-defined kinetic rate laws, such as mass action or enzymatic kinetics. In ChemChaste, both the stoichiometry and kinetic rates of these reactions are defined through a file-based user interface (see next section and SI, section S2.2.2). Furthermore, bulk reactions can be assigned to a specific sub-domain (of the `Domain Field)` of the mesh. To assist with the assignment of kinetic laws to reactions, ChemChaste implements specific classes describing different kinetic laws. In ChemChaste, chemical species may be provided with a set of properties: name, diffusivity, mass, valence, Gibbs formation free energy. These properties can be linked to affect the rate of diffusion or rate of a given reaction within which the species participate. Furthermore, when the `Domain Field` contains sub-domains, the domain varying chemicals’ properties may be stored in upstream inheritance classes. This allows simulating changes in diffusivity due to spatial heterogeneities (e.g. bulk media vs. biofilm or tissue). Within the ChemChaste code, these chemical associated parameters can be called by the PDE diffusion functions or reaction systems for the correct sub-domain.

### 2.4 File-based user interface

ChemChaste introduces a file-based interface to enable its use by a wider audience. In particular, ChemChaste has two main user-interface systems, one to provide the `Domain Field` and diffusion properties and one for defining the `Reaction System`, which together characterise a heterogeneous reaction-diffusion model. The `Domain Field` files contain the information required to produce the FE mesh and define the labelled sub-domains. This file also defines any varying BCs and/or diffusion rates for bulk chemicals. The user supplies a comma separated values (CSV) file of labels denoting the sub-domains and a text file of the associated label keys (see SI, section S2.2 for an exemplar `Domain Field` file). Further CSV files of initial species values, boundary conditions, and diffusion rates on a sub-domain basis may also be specified. These files fully characterise the conditions of the simulation space, while the reaction dynamics are detailed in a separate reaction file.

The `Reaction System` file encodes the bulk, cellular, and coupling (i.e. membrane and transport) reactions as described above. For the bulk reactions each sub-domain can have an associated, separate reaction system file. Another file is used to define the cellular reaction system. Within this cell file, coupling reactions are defined with at most one membrane reaction file and one transport reaction file, each containing a set of reactions of the respective type. All reaction files follow a set format; name of reaction kinetics, chemical equation involving the species, then the kinetic parameters used by the rate laws (see SI, section S2.2.2). Further rate laws may be implemented by the user, which will then be utilised in the same way as the supplied rate laws (see SI, sections S4–S6 for details). Overall, the information stored within these files is sufficient to select the desired reaction class, formulate reaction terms and implement concentration changes when solved within the simulation.

## 3 Results

ChemChaste presents a hybrid continuum-discrete modelling framework for the simulation of individual cells within a chemically active environment. As shown in Figure 1 and discussed in the Methods section, the framework is composed of an array of different modules building upon each other to fulfil the simulation needs. Here, we verify and demonstrate the functionality of ChemChaste by considering each of these key modules in turn. The accuracy of the PDE solvers was tested through solving the Fisher-Kolmogorov-Petrovsky-Piskunov (Fisher-KPP) equation showing a strong agreement with an analytic series expansion (Section 3.1). The simulation of multiple PDEs using the ChemChaste reaction system and file interface system was demonstrated through producing diffusion-driven spatial patterning and temporal oscillations of the Schnakenberg reaction system (Section 3.2). Finally, an exemplar coupled cell simulation was implemented involving a cooperator-cheater system based on enzyme excretion (Section 3.3).

### 3.1 Spatial simulation accuracy in ChemChaste: Fisher-KPP equation

To verify and demonstrate the PDE solving capabilities in ChemChaste, a single PDE with a known analytical solution was implemented. The chosen system was the Fisher-KPP equation, which has been used to model the propagation of an invasive species through a population (Fisher, 1937; Murray, 2002) and admits travelling wave solutions with an analytically resolved minimum wave velocity (El-Hachem *et al*., 2019). The corresponding reaction-diffusion equation includes a logistic growth source term,

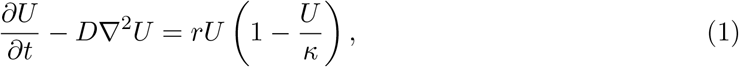

where *U*(x, *t*) ≥ 0 is the size of the invasive species population at position x = (*x, y*) and time *t*, and the positive parameters *D, r* and *κ* denote the diffusion coefficient, linear growth rate and carrying capacity of the invasive species, respectively. For suitable initial conditions, it is known that this system exhibits pulled travelling wave solutions of the form *U*(*z*) where *z* = *x* – *ct* and *c* ≥ 0 is the wave velocity. It can be shown analytically that the front of these waves travels with a minimum velocity defined by

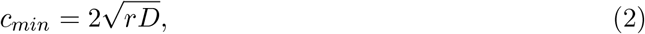

while the observed velocity, *c* ≥ *c_min_*, is dependent on the initial conditions (Murray, 2002; El-Hachem *et al*., 2019).

We implemented the Fisher-KPP equation in a ChemChaste simulation using equation (1) and setting the parameters to unity {*D*, *r*, *κ*} = 1. We considered a rectangular bounded domain Ω ∈ [0, 10] × [0, 100] and impose zero-flux boundary conditions (BCs) and record a 1-dimensional slice across the domain. The simulations were initialised with a strip of invasive species bordering the left boundary of the domain, 0 < *x* < 1:

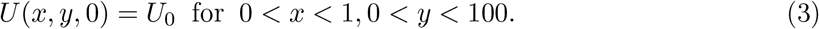

For equation (1) the minimum wave speed with the selected parameter set is given by *c_min_* = 2.

The FE methods within ChemChaste were used to solve equation (1) subject to the boundary and initial conditions. A travelling wave solution was identified across the one-dimensional domain slice and compared to the analytical solution of the one-dimensional Fisher-KPP equation (Loyinmi and Akinfe, 2020), given by

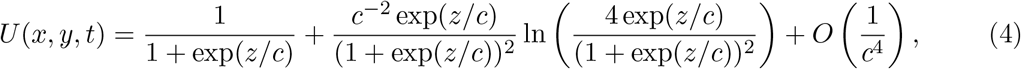

where *z* = *x* – *ct* denotes the travelling wave coordinate.

The results were visualised using ParaView (Ahrens *et al*., 2005). Two tests were considered: comparing the travelling wave front solution produced by the ChemChaste simulation vs. the analytic form given by equation (4), and comparing the simulations’ convergence stability under decreasing temporal and spatial step size. Results for both tests are given in Figure 2, and show a good agreement between the ChemChaste simulation output and expected results determined through analytic solutions. Additionally, the convergence with decreasing temporal and spatial step sizes suggest stable numerics albeit with the waves showing longer accelerating phases than the expected analytic top-hat gradient. Therefore the ChemChaste implementation was able to correctly simulate dynamics (in this case, the travelling wave phenomenon) in simple PDE with stable and accurate numerics.

**Figure 2:**
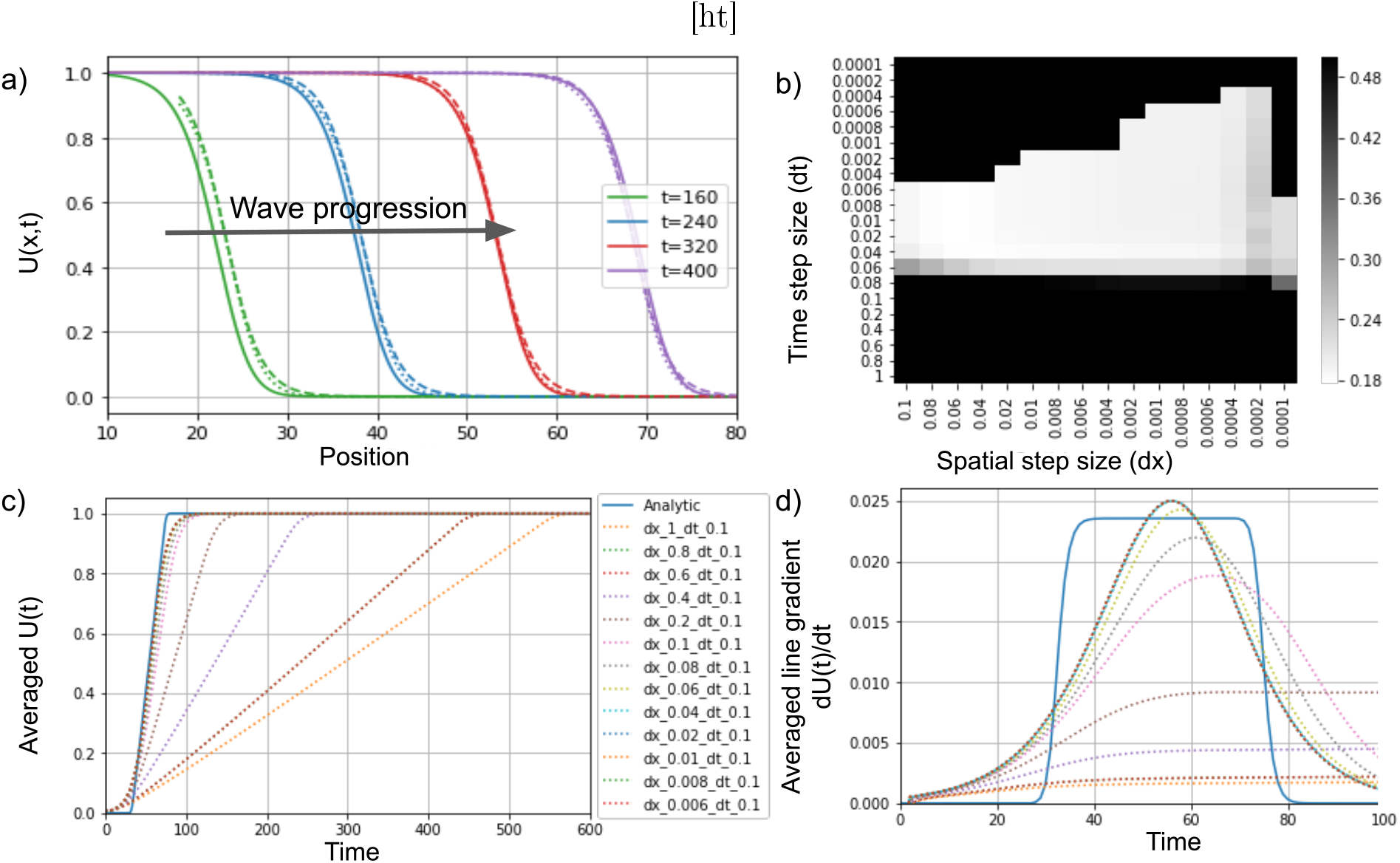
ChemChaste simulations of the Fisher-KPP equation. a) Plot showing the progression of an expanding wavefront through the domain (solid line). The simulation results are accompanied by the analytic solution for the zeroth (dashed line) and first order (dotted line) expansion in terms of 1/*c*^2^ in equation (4). The wave speed in simulation is initially faster than the analytical minimum wave speed *c_min_* = 2, calculated with equation (2), but with agreement at later times implying the correct asymptotic wave velocity has been reached. b) Heat-map of *L*^2^ convergence scores for simulations using a range of spatial and temporal step sizes. The simulations for given step sizes are compared to the analytically determined value with the lower scores suggesting closer values. A threshold was utilised reducing higher scores to 0.5. This includes simulations whose numerics diverged. c) Traces for the solutions U(t) averaged across the domain for different spatial and temporal step sizes. The traces converge to the analytical solution with decreasing step size. d) The gradients of the slopes in plot c) sharing the same legend. The gradients are suggestive of the velocity of the wave passing through the domain.

### 3.2 Modelling multiple, diffusing and reacting chemicals in ChemChaste: Schnakenberg reaction-diffusion system

ChemChaste builds upon Chaste’s PDE solvers to enable the simulation of multiple PDEs over the domain. While Chaste is restricted to solving three PDEs, ChemChaste’s limiting factor is solely the available computational resources. To test the multi-dimensional PDE simulation, and to verify the file interface system, we implemented the well-studied two species reaction system commonly known as the Schnakenberg system (Li *et al*., 2018; Schnakenberg, 1979) and shown in equations (5)–(7). When these reactions are modelled with mass action kinetics they are shown to display temporal oscillations and diffusion driven spatial patterning for distinct, defined parameter regimes (Al Noufaey, 2018; Murray, 2003). These phenomena were reproduced here using ChemChaste.

The Schnakenberg reaction system involves two chemical species *U*, *V* which are produced, inter-converted, and removed via the reactions

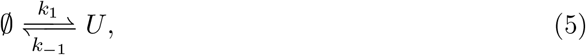

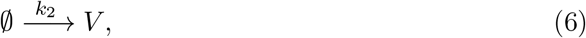

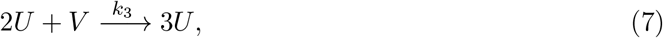

where the reaction rate constants parameters are denoted by *k*_1_, *k*_−1_, *k*_2_, *k*_3_. Applying mass action kinetics to these reactions yields the reaction ODEs

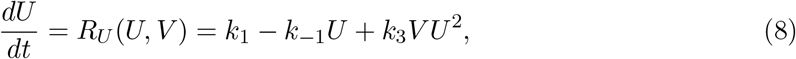

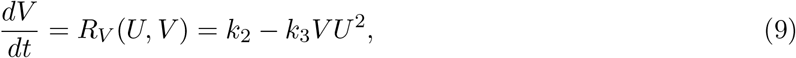

where the reaction rates *R_U_, R_V_* describe the change of each species’ concentration in a given timestep and also provide the source terms to the reaction-diffusion PDEs. The PDEs are satisfied across the whole two-dimensional domain space, Ω, and are given by

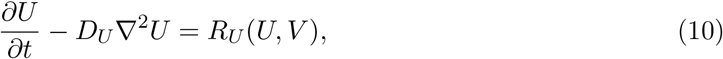

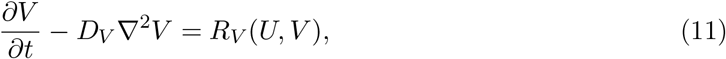

where *D_U_*, *D_V_* are the spatially homogeneous isotropic diffusion coefficients. Here, we consider a square bounded domain Ω ∈ [0, 100] × [0, 100] which are subject to zero-flux Neumann BCs

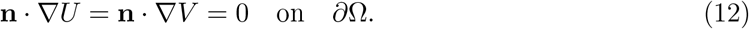

Each simulation begins with the randomly perturbed initial conditions defined on each node of the FE mesh,

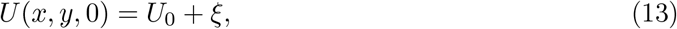

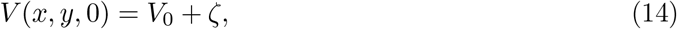

where *ξ*, *ζ* ~ *Uniform*(−1, 1) are uniformly distributed random fields bounded by the interval [−1, 1].

Two parameter sets were considered: one for temporal oscillations; and one for diffusion-driven patterning (Al Noufaey, 2018). Temporal oscillations are present when the homogeneous system, equations (8)–(9), display limit cycle behaviour. Spatial patterning across the domain occurs when the spatially uniform steady-state solution to equations (10)–(11) is linearly stable in the absence of diffusion (*D_U_* = *D_V_* = 0), but linearly unstable in the presence of diffusion. The resultant spatial patterning in the 2D concentration maps are referred to as displaying diffusion-driven instabilities (DDI) or Turing instabilities (Murray, 2003; Turing, 1952; Page *et al*., 2003; Maini *et al*., 1992). These dynamical cases were found to occur for specific parameter sets, as listed in Table 1.

**Table 1:** Parameters used in the Schnakenberg reaction simulation. The values were selected based on analytical solutions of this system and to demonstrate the possible oscillatory and patterning dynamics.

| Case | $k_1$ | $k_{-1}$ | $k_2$ | $k_3$ | $D_U$ | $D_V$ | $U_0$ | $V_0$ |
| --- | --- | --- | --- | --- | --- | --- | --- | --- |
| Figure 3a: Oscillations | 0.5 | 2.2 | 1.5 | 1.0 | 0.5 | 0.5 | 0.91 | 1.67 |
| Figure 3b: Patterning | 0.1 | 1.0 | 0.9 | 1.0 | 1 | 40 | 1.0 | 1.0 |

These parameters were determined through considering small linear perturbations for conditions which provided the expected phenomena in the two cases, equations (8)–(9) and (10)–(11), and selecting parameter sets which satisfy the algebraic equations (Murray, 2003), (see SI, section S3 for details). The values *U*_0_, *V*_0_ were used as the initial conditions for the two cases.

We have verified, using ChemChaste, that this model exhibits the expected spatio-temporal dynamics for the tested parameter regimes (see Figure 3). These results are as expected for the parameters used, based on analysis of equations (8)–(9) and (10)–(11). Therefore these tests verify that ChemChaste was able to both correctly parse the chemical reaction files and simulate multi-chemical reaction-diffusion systems capable of complex dynamics and patterning.

**Figure 3:**
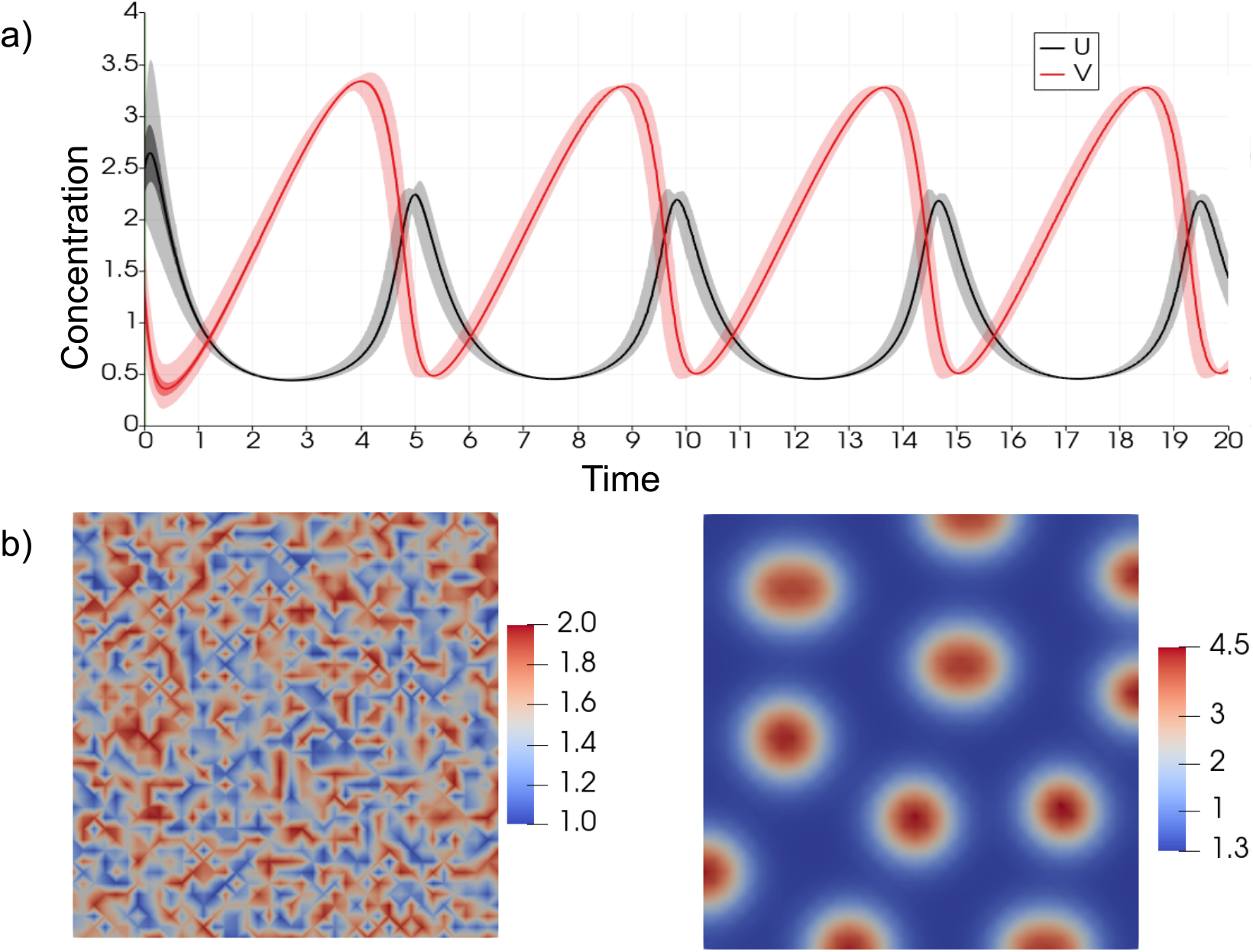
The Schnakenberg reaction system showing the oscillatory and patterning dynamics. a) The two curves show the concentration of U and V averaged over the nodes in the domain for each time step from the simulations run using oscillatory regime parameters. b) Domain maps of the initial and final (i.e. steady state) distribution of U and V in simulations using parameters for the patterning dynamics (see Table 1).

### 3.3 Coupled cell-chemical environment simulations in ChemChaste

A main motivation behind developing ChemChaste was to simulate a hybrid continuum-discrete model of cells within a chemically reactive environment, where bulk and cell-secreted chemicals and other entities such as proteins can diffuse as well as react. This is a common biological scenario, as seen for example in the case of microbial utilisation of cellulose or other complex resources, which must be treated by enzymes before a cell can metabolise or uptake them (Flint *et al*., 2012). The core aspects of this scenario, i.e. a cell-secreted enzyme mediating a reaction in the bulk is also found in cases outside of substrate uptake, for example in de-toxification of the environment (Zerfass *et al*., 2019). In ChemChaste, this scenario is readily modelled through implementation of bulk reactions and coupling of cellular metabolic reactions and environmental PDEs.

Here, we provide a simplistic, toy example for illustrative purposes and for testing ChemChaste implementation of cellular reactions and cell-environment coupling. More detailed and realistic simulations can be readily constructed by users, through developed ChemChaste user interface. For the exemplar test case, we modelled a growing cell population harbouring two cell types, along with a chemical resource (i.e. substrate) that is not readily taken up. One cell type - termed cooperator - excretes an enzyme that can allow the internalisation of the substrate, while the other cell type - termed cheater - does not excrete the enzyme but can also internalise the enzyme-bound substrate (Figure 4a). The cells process the internalised substrate to produce a pseudo chemical species (called ‘biomass’), which is used as a proxy for monitoring cell growth. Once the cellular biomass concentration reaches a threshold value the cell divides into two, the parent and offspring, sharing the internal concentrations equally between both parent and offspring cell. The offspring cell is placed at a random neighbouring location around the parent cell and the population undergoes positional updating to accommodate the new cell.

**Figure 4:**
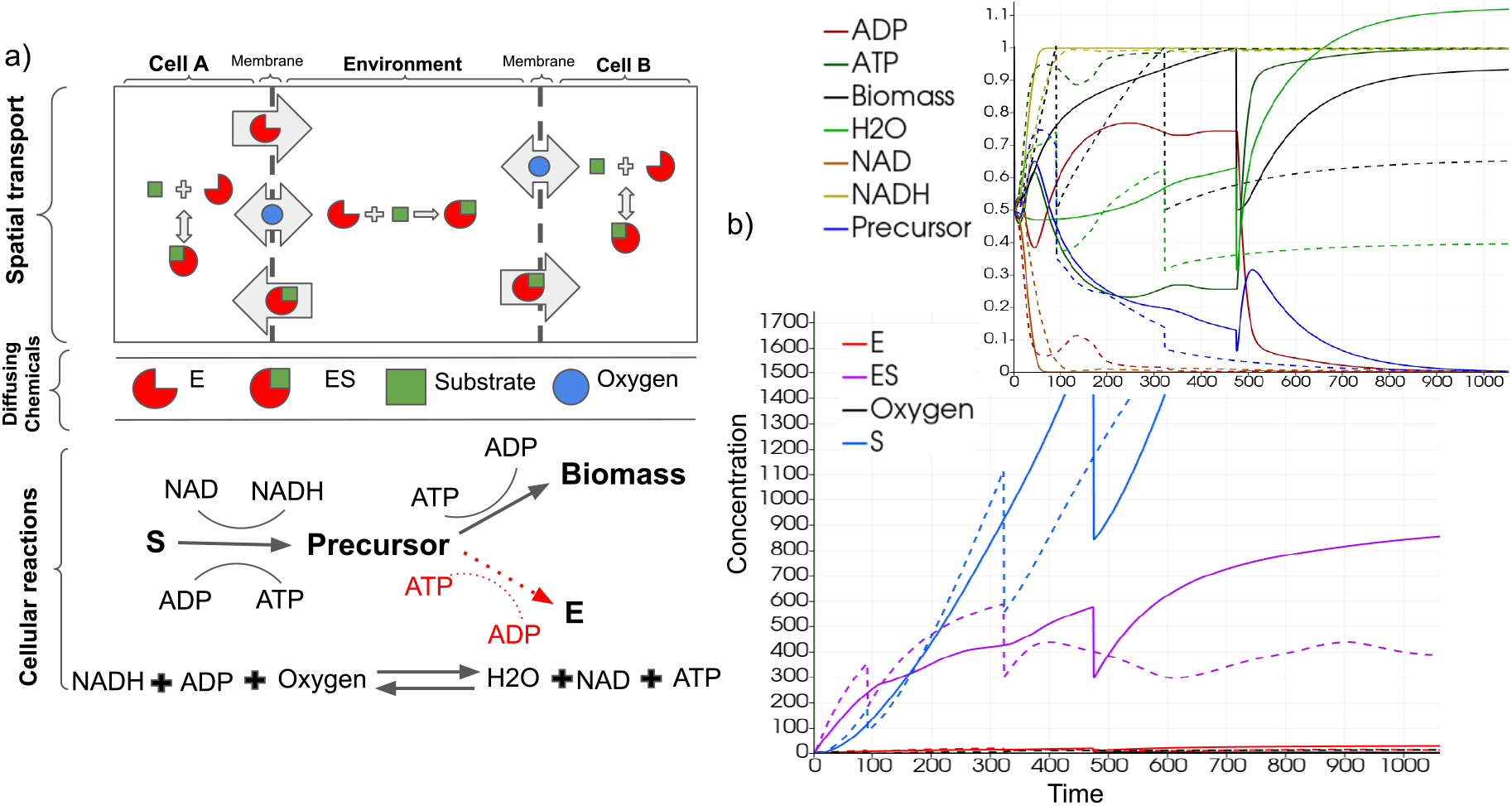
The simulation schematic and results for the exemplar cellular model with cell-environment coupling. a) Cartoon showing the two cell types and cellular reaction system implemented in the simulations. One cell type, the ’cooperator’, excretes an enzyme that can bind an environmental substrate, while the other - the ’cheater’ - does not produce the enzyme (top). Both cell types can take up the enzyme-substrate complex and process it through a series of internal reactions (bottom). Note that the enzyme producing pathway is only active in the cooperator cells, which has to invest substrate between this pathway and biomass producing pathway. b) The concentrations for each chemical within the cell are displayed over time for a cell of both types; cooperator (solid) and cheater (dashed) lines. The main plot shows the concentrations of ES and S (chemicals harvested from the environment). The inset shows the concentrations of the cell-internal chemicals. Sharp changes in cellular concentrations are due to cell division and sharing of chemicals between the parent and offspring.

Previous agent-based simulations of growing cell populations harbouring cheater and cooperator types have found spatial segregation of cell types within the population (Nadell *et al*., 2010; Mitri *et al*., 2016; Momeni *et al*., 013a). This cell sorting is linked to the disparity in growth rates of the two species, which may be due to substrate availability and dependency, and is of interest in game theoretic investigations of mutual interactions in biofilms (Tudge *et al*., 2016; Rubin and Doebeli, 2017). The presented simulations are conceptually similar to these previous studies, but differ in their mechanistic implementation of substrate scavenging, as a cooperative trait, as well as the inclusion of both substrate and oxygen diffusion in the bulk.

In the presented model the two types of cells were introduced into the simulation domain which contains two chemicals which diffuse in the bulk; oxygen (*O*_2_) and a substrate, *S*. Furthermore the cells excrete and take up a scavenging enzyme, *E*, the enzyme-substrate complex, *ES*, and *O*_2_, which freely diffuses in the bulk. To capture dynamics of cell growth, a simple metabolic network is implemented in each cell, defined by the following toy reactions that abstract biomass generation and the main respiratory and fermentative metabolic pathways:

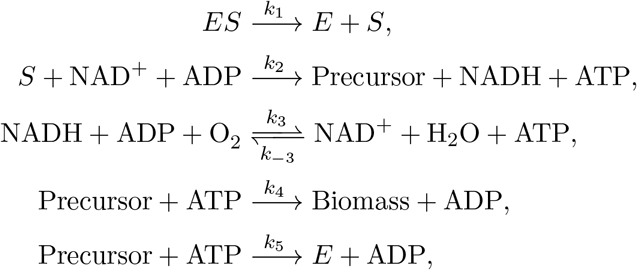

where NAD^+^, NADH, ADP, and ATP are the usual energy and electron carrier molecules internal to the cell. These toy reaction set captures substrate uptake (reaction 1), re-cycling of NAD^+^/NADH and ADP/ATP pairs through fermentative and respiratory pathways (reactions 2 and 3), and biomass and scavenging enzyme production through ATP investment (reactions 4 and 5). For the simulations, these reactions are modelled with mass action kinetics with shown reaction rate constants. All reaction rate constants were set to 1 in both cell types, except for *k*_5_, which is set to zero in the cheater cell type. The overall simulation schematic for this cellular system is shown in Figure 4.

In addition to the cellular reaction network, we implemented bulk reactions for the enzyme binding to the substrate in the extracellular media, the enzyme being degraded in the bulk, and the diffusion of the substrate (S), enzyme (E) and the enzyme-substrate (ES) complex.

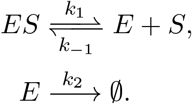

The parameters for these reactions were scaled for computational efficiency and are given in SI, section S2.3. We performed simulations through the hybrid continuum-discrete solvers introduced in ChemChaste. A reaction-diffusion PDE was solved over the domain for the diffusing species {*E, S, ES, O*_2_} with Neumann BCs at the domain boundary. The Neumann boundary conditions allow continual replenishment of substrate to drive the system. The cells were placed in the centre of this domain with a single cell of each type, and allowed to grow over the simulation course, as shown in Figure 5. The chemical concentrations in each cell and the bulk were recorded over the simulation. Note that initial substrate levels at the beginning of the simulation are low, but will linearly increase due to the implementation of the Neumann boundary conditions. Additional boundary conditions, like Dirichlet type, can be defined per the user files.

**Figure 5:**
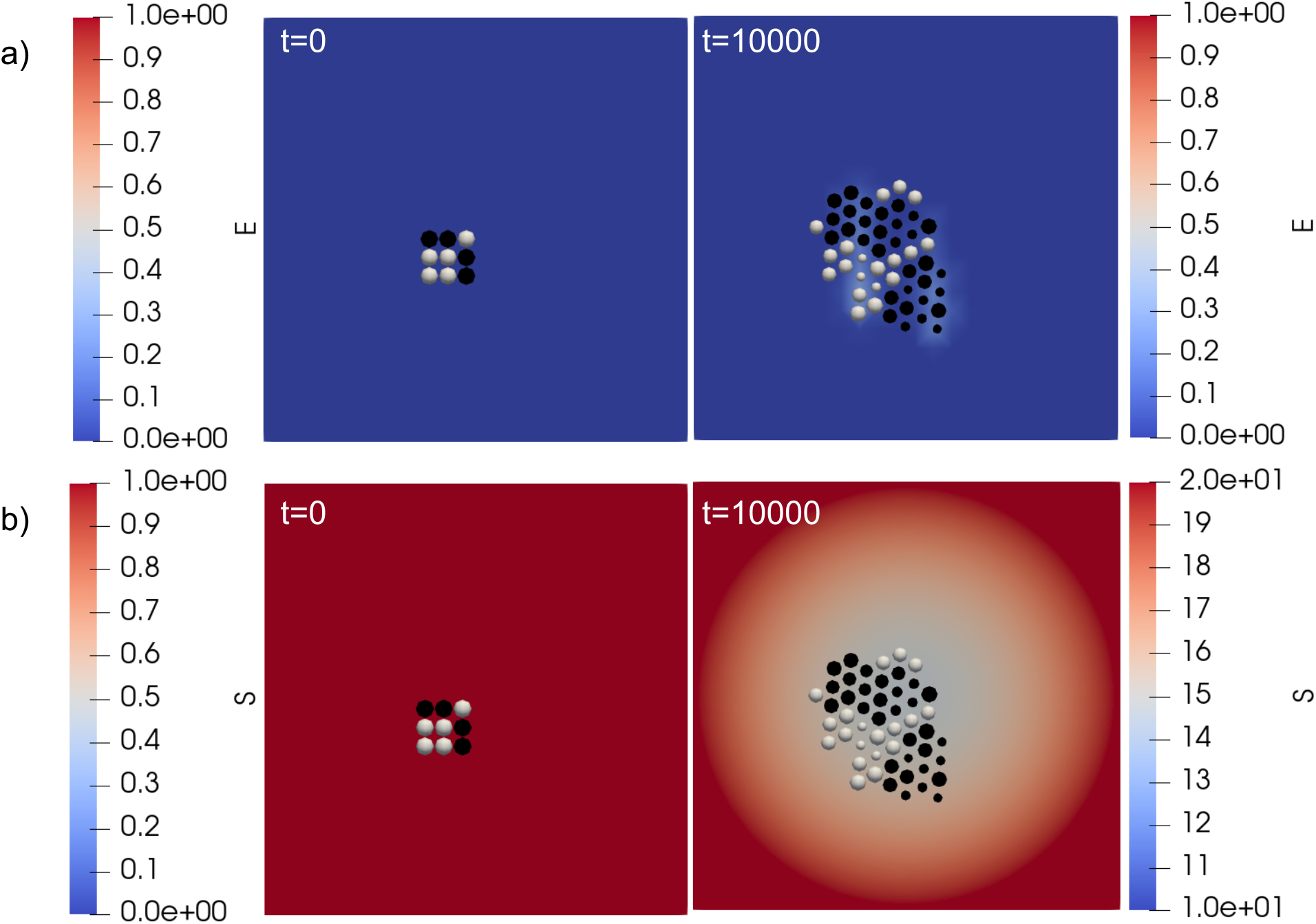
Domain maps showing the cells and the spatial concentration of the enzyme and nutrient species, *E* and *S*, in the bulk at the towards the beginning, *t* = 0 and *t* = 100, and at the end, *t* = 10000, of the simulation. The cooperator and cheater cells are coloured white and black, respectively. The upper row (a) shows simulation results with the enzyme (E) concentration plotted across the domain. The lower row (b) shows simulation results for the substrate (S) across the domain with the replenishment of the substrate at the boundary.

We show the dynamics of cellular and bulk chemicals in Figure 4 and 5. While Figure 4 is focused on the cell concentrations, Figure 5 demonstrates the impact that the cells have on local chemical concentrations. In Figure 5a, we see a higher enzyme concentrations in the vicinity of cooperator cells. This is as expected, since these are the cells excreting the enzyme. We expect that such higher local concentrations of enzyme will be enhanced with lower enzyme diffusion rates and enzyme degradation rate in the bulk. In Figure 5b, we see the substrate concentration, with higher values at the domain edge (due to influx of substrate) and lower values near the cell population (due to cellular uptake). Evaluating Figures 4 and 5, together, we see a greater uptake of the substrate by the cooperator cells and a greater rate of cell biomass increase, compared to the cheater cells. Thus, the localised pockets of high enzyme concentrations around cooperator cells can lead to their growth rate surpassing that of cheaters and subsequently lead to a spatial segregation of the two cell types. While further simulations with different parameter sets are needed to fully confirm these dynamics, the presented results provide an exemplar implementation of cellular simulations in ChemChaste and confirm expected cooperator-cheater dynamics.

We conclude that the presented toy model and exemplar implementation of a cellular simulation demonstrate ChemChaste’s flexibility and capabilities in developing models featuring cell-environment coupling along with environmental reaction-diffusion.

## 4 Conclusion

We have presented ChemChaste, a computational framework for hybrid continuum-discrete modelling of multi-cellular populations coupled to chemical reaction-diffusion systems. In contrast to existing computational frameworks, ChemChaste facilitates chemical couplings between bulk and cellular metabolic processes through an arbitrary number of diffusing chemicals that can undergo chemical reactions in the bulk and that can have spatially heterogeneous diffusion coefficients. ChemChaste simulations are implemented using a simple file-based interface and can be used to implement different biological and chemical scenarios for modelling complex cell-environment chemical coupling and resulting emergent phenomena.

We have presented several exemplar simulations in ChemChaste, which produce the expected dynamical behaviours in given parameter regimes. These exemplars were specifically chosen to demonstrate ChemChaste’s functionality and flexibility, instead of presenting an exhaustive list of the possible phenomena that may be investigated using this tool. Applications of immediate interest can include different observed cases involving coupling between cellular physiology, cell excretions, and environmentally diffusing reactions such as metabolic switching of cell types coupled to a reactive environment (Ratzke and Gore, 2018; Varahan *et al*., 2019), coupled chemical reactions in the bulk and within cells (Turing, 1952), coupling between cell secreted enzymes, signalling, and motility (Weijer, 2009), and cell-chemical systems presenting spatially varying diffusion coefficients (e.g. within and outside of a tissue) (Liu *et al*., 2015).

Some of these investigations may require further expansion of ChemChaste. Such as the extension into 3D modelling which would require expanding the ChemChaste code. However, for users proficient in C++ the addition of new classes is straightforward through the addition of new user-defined classes to the ChemChaste C++ class hierarchy utilising the modular structure of the framework. In this way we hope ChemChaste will prove a useful tool for investigating the chemical mechanisms behind a range of phenomena in spatially organised biological systems.

## Acknowledgements

The authors thank Aydar Uatay for useful discussions about Chaste.

## Supporting information

### S1 Brief introduction to the finite element method and the hybrid continuum-discrete model used in ChemChaste

At the heart of ChemChaste lies a suite of finite element (FE) solvers, which are used to numerically solve systems of reaction-diffusion partial differential equations (PDEs). These equations track the spatiotemporal dynamics of a set *C* of chemical species over a bounded rectangular domain 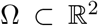 whose boundary is denoted ∂Ω. Each chemical species *c* ∈ *C* is associated with a scalar concentration field, with the associated state variable 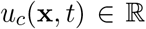 denoting the concentration of the chemical at position x ∈ Ω at time *t*.

ChemChaste couples the reaction-diffusion system to an agent-based cell system to model a spatially distributed cell population. We define a set of cells, *p* ∈ *P*, within the domain which are modelled as point sources at position x*_p_* ∈ Ω. These cells perform reactions independently of the domain reactions and exchange chemical concentrations with the domain. These exchanges are described by the transport law 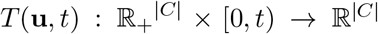 controlling the chemical concentrations passing between the bulk and the cell.

During a simulation the vector of state variables, 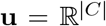, evolves through the parabolic PDE system

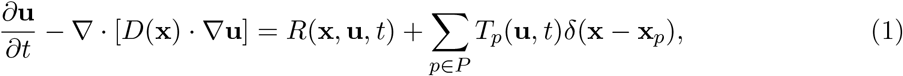

where *T_p_*(**u**, *t*) is the source/sink contribution by the cell *p* located at position x*_p_*, 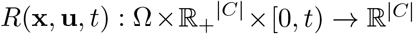 is the reaction ordinary differential equation (ODE) system defined over the domain, and *δ* is the Dirac delta distribution. ChemChaste models the spatially discontinuous distribution by a top-hat distribution with maximum value 1 and 2D extent covering the Gauss point associated with location x (see section S1.2). We solve equation (1) as an initial boundary value problem (IBVP) with given initial conditions (ICs) and boundary conditions (BCs).

For each chemical species, we allow for either fixed-value Dirichlet BCs of the form

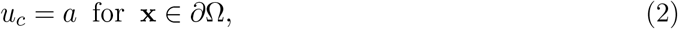

or fixed-flux Neumann BCs of the form

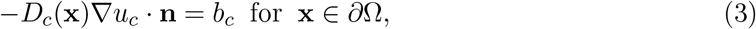

where **n** is the unit outward normal, *D_c_*(x) is the (*local*) diffusion coefficient of chemical species *c*, and 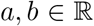 are constants. Isolated conditions, *b_c_* = 0, are assumed for chemicals without a user defined Neumann BC.

While the BCs are defined on the whole boundary ChemChaste handles chemical initial conditions through labelling regions of the domain. A set of regions, *S*, are provided using the file based input (see section S2.2), which link to chemical concentrations. Let *s* ∈ *S* label a region Ω*_s_* ⊆ Ω asssociated with an initial chemical concentration vector 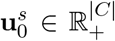. For each position x ∈ Ω, the initial conditions are implemented with sharp discontinuous boundaries

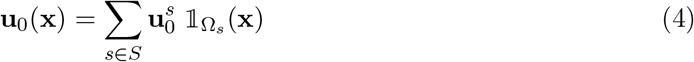

where

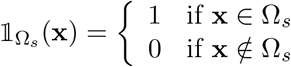

is the indicator function.

We solve the above system numerically using a FE method, which divides the spatial domain Ω into a discrete mesh comprising nodes and elements, and converts the PDE problem into a coupled system of algebraic equations (Pathmanathan, 2012).

#### S1.1 Finite element method for a reaction-diffusion PDE coupled to a cell mesh

To apply the FE method to the PDE system (1), we first derive the weak formulation of the problem. We start by discretizing time into discrete timesteps *t^m^* = *m*Δ*t* and applying a semi-implicit scheme to the time derivative in (1), obtaining

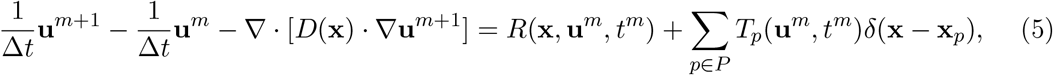

where **u**^*m*^ denotes the approximation to the chemical concentration vector **u**(x, *t^m^*) for x ∈ Ω. Next, we multiply by a vector **v** of arbitrary ‘test functions’ with each element *v* ∈ **v** defined in a function subspace of the Sobolev space 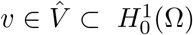 and integrate over the domain Ω. Rearranging, we obtain

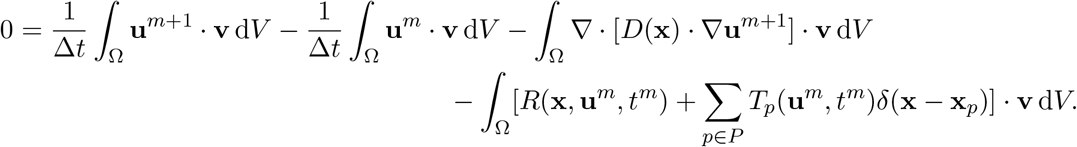

Writing the diffusion term as 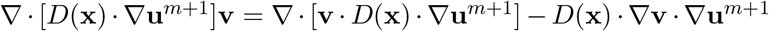, and applying the divergence theorem, we obtain

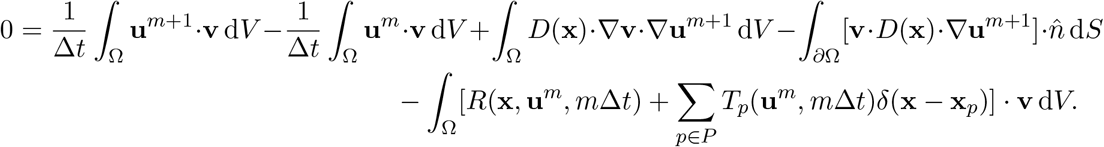

We then consider the BCs as *v* = 0, ∀*v* ∈ **v** by the natural boundary definition for Dirichlet types and 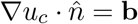 for the Neumann types to get

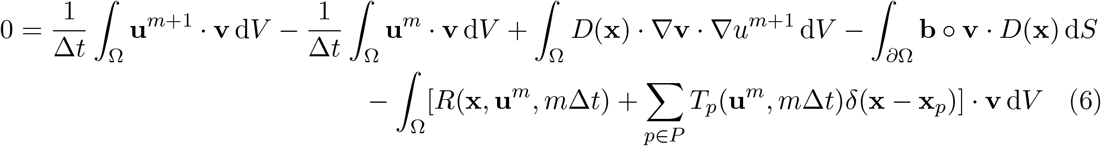

which defines the weak formulation of the PDE as a variational problem, with ∘ representing the element-wise product. This allow us to determine functions **v** in order to find the next state variable vector **u**^*m*+1^ given the previous **u**^*m*^ state variable vector such that the remaining Dirichlet BCs for **u**^*m*+1^ are satisfied (Logg *et al*., 2012). This problem is solved on the nodes of a discretised spatial domain, i.e the FE mesh nodes.

To implement the FE method a mesh is placed over the domain, Ω. This mesh provides a space of discrete nodes and for each node *i* at position *n_i_* on the domain Ω \ ∂Ω, i.e each node not on the boundary, a set of linear basis functions *ϕ_i_* are defined {*ϕ*_1_, *ϕ*_2_, …, Φ*_N_*} where *N* is the number of nodes in the domain. We solve equation (6) on the discrete node points and interpolate onto a fine rectilinear grid to approximate the domain locations x. We refer to these interpolated locations as Gauss points and discuss the interpolation in Section S1.2. The Gauss point x is given by the product of the node location **n***_j_* and the linear test basis function *ϕ_j_*;

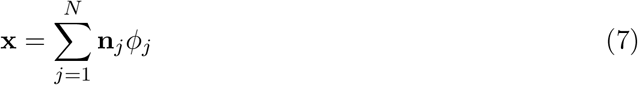

Using the mesh interpolation the trial function can be approximated by

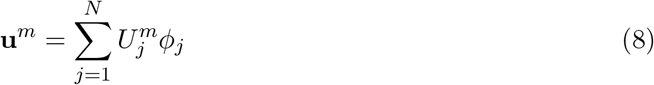

where **U**_*j*_ is the nodal value. Additionally, we find the gradient of the trial function is given by

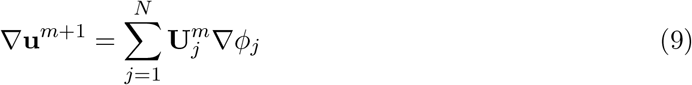

We are free to restrict the basis functions, *ϕ_i_*, to functions that satisfy the requirements for the test functions over the Ω. Therefore we may substitute ***ϕ**_i_* for **v**

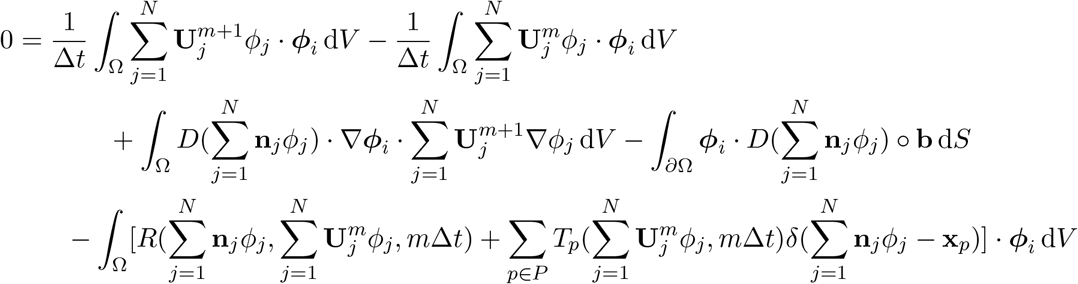

which can be converted into the algebraic system

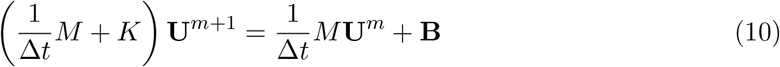

by defining *K*, *M* and **B**, as

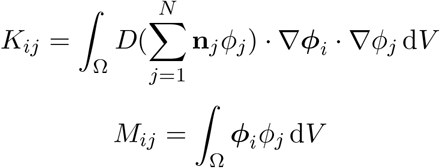

and

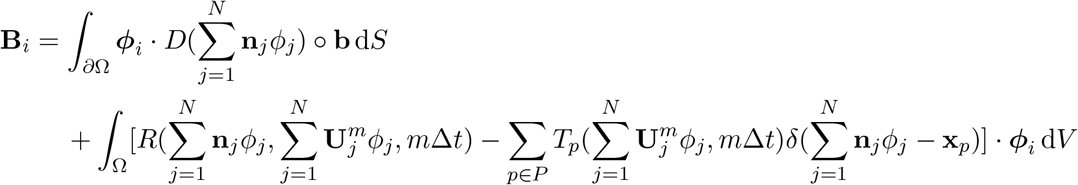

Dirichlet BCs are applied by altering the matrix, *K*, and vector, *B_i_*, for the contributions of nodes on ∂Ω while Neumann BCs are contained within the definition of *B_i_*. The coupled cell system is performed by solving a system of ODEs and agent properties for the cells and solving the linear system (Equation (10)).

#### S1.2 Interpolation of the nodal solution to Gauss points and coupling to cells

As motivated in Section S1.1, the FE method employed discretises the domain space using elements formed by nodes, solves a linear algebraic system on those nodes and then interpolates the solution within each element. While Chaste has been developed to run simulations in 1, 2, or 3 spatial dimensions, the current version of ChemChaste has been restricted to simulate 2-dimensional domains. Chaste utilises linear Lagrange elements for creating the FE mesh (Logg *et al*., 2012). To construct the FE mesh for a 2-dimensional domain the domain is partitioned into a finite set of triangles, **T**, which cover the space;

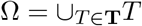

Each triangle contains a triplet of nodes where a pair of nodes may be shared by adjacent triangles, shown in Figure S1a. The mesh is defined by the set of *N* nodes, *L* = {*l*_1_, *l*_2_, …, *l_N_*}.

This triangulation process and the resulting elements define a so-called Sobolev space, and permits the choice of a linear basis, *ϕ*, for the test functions (Shapira, 2012). In Chaste, the linear Lagrange basis is used such that for node triplet at triangle locations 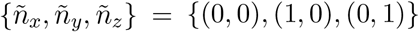 is spanned by a basis of the form;

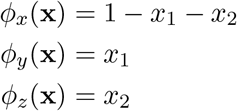

where the test function is approximated by the linear superposition of the values at the nodes

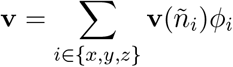

The positions within the triangle are discretised by a fine grid for computational purposes. These grid locations 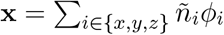 are known as the Gauss points. The local points are mapped to global locations as shown in Figure S1a. ChemChaste introduces the cell contribution to equation (10) if a cell is located at the Gauss point, Figure S1b.

**Figure S1:**
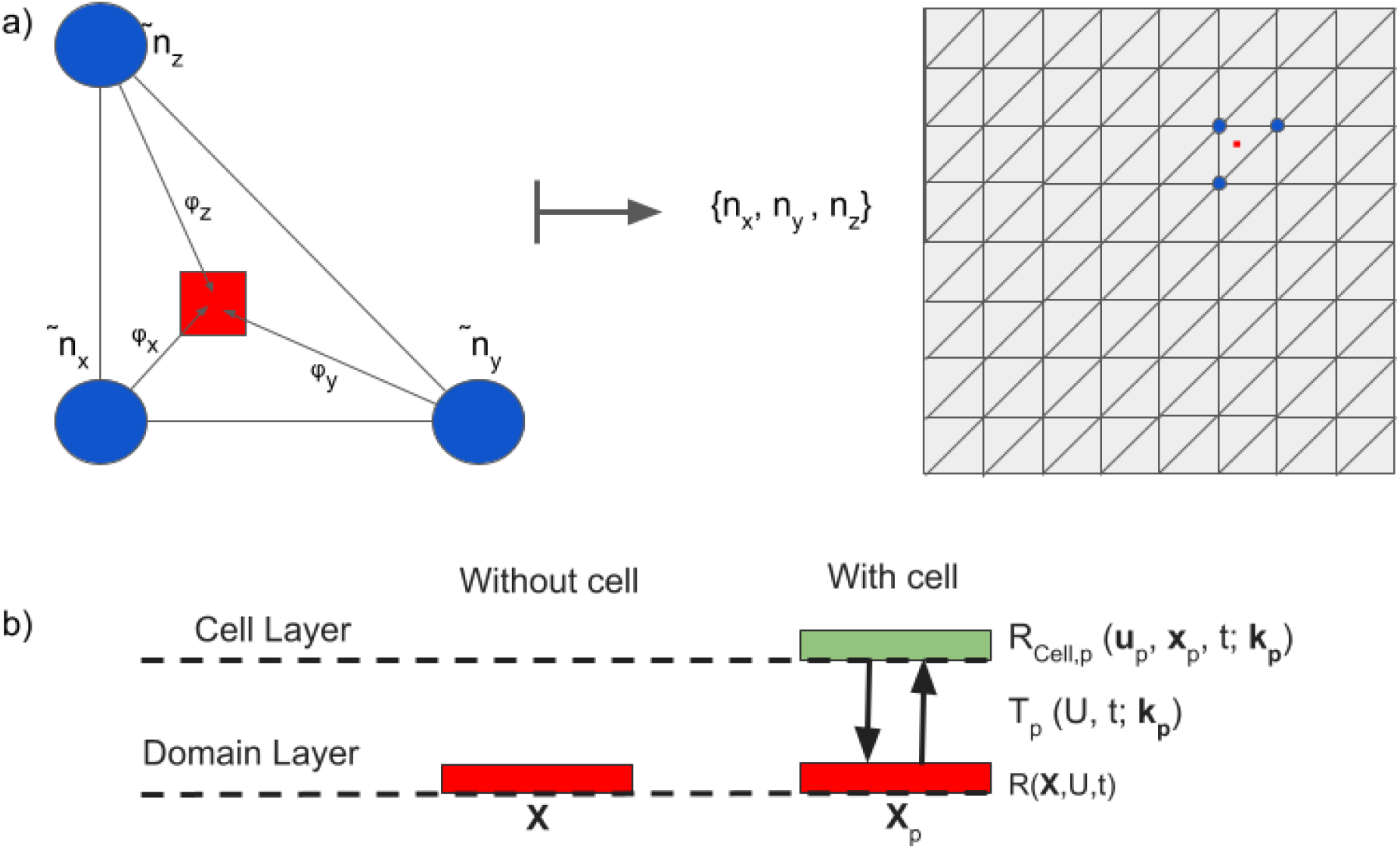
Cartoon showing the interpolation procedure from finite triangle elements to Gauss point and coupling to cell agents. a) The nodes (blue) of the triangle are located at positions 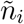 and each node contributes to the particular Gauss point (red square) through the node’s basis function *ϕ_i_* for node *i* ∈ {*x, y, z*}. The node locations and other interpolated quantities are mapped from the local reference triangle to the global mesh i.e by applying a mapping function to the node position 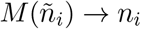, which may, in general, rotate and stretch the reference triangle. b) Cells have a point location which share the location of a Gauss point. Each point in Ω is associated with a spatially dependent reaction system, *R*(x, *U*, *t*) and may also be associated with a cell. These cells (green) contain their own reaction system, *R_Cell,p_*(**u**_*p*_, x_*p*_, *t*; **k** – *p*), and are coupled to the domain through a transport law *T_p_*(*U*, *t*; **k**_*p*_).

#### S1.3 Simulations of chemical reaction PDEs without cells: Chemical reactions *R*(u)

Reaction-diffusion simulations in ChemChaste are created from a set of user-defined chemical reactions as provided in configuration files. Let the set of all chemical species within a system be denoted by *C* and the concentration of a particular species *c* ∈ *C* be denoted *u_c_* where 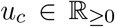. For a given reaction, let 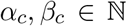 denote the unsigned stoichiometry of species *c*. *β* >= 1 if the species is a product of the reaction, where *β_c_* is the number of the species produced per one instance of the reaction. Conversely, *α_c_* is the number of the species consumed within the reaction; *α* >= 1 if the species is a substrate species of the reaction. The change in concentration, hence value for *u_c_*, is given by

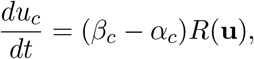

where the function *R*(**u**) defines the reaction dynamics (Table S1).

For a given reaction, let *S* ⊆ *C* denote the set of substrates in the reaction, *P* ⊆ *C* the set of products, *kf* the forward reaction rate constant, *kr* the reverse reaction rate constant, and let 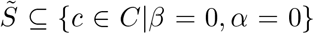 be the set of spectator species. The spectator species affect the reaction rate while remaining unchanged themselves. As an examplar, consider a set of chemicals in the domain *C* = {*A, B, C, D, E, F*} and a chemical reaction involving two substrate species *S* = {*A, B*}, two products *P* = {*C, D*} and a single spectator species 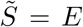. The chemical reaction may be written as;

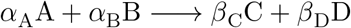

where the change in concentrations will depend on *R*(**u**);

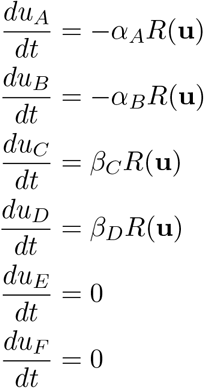

*E* is a spectator species and therefore is not produced or consumed in the reactions but affects the value for *R*(**u**) and 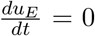 as *E* as a steady state for *E* is assumed within the context of this reaction system. Implicitly,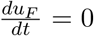 as *F* as chemical *F* is present within the domain but is not involved within the reaction system.

Table S1 presents the pre-implemented reaction rate laws for bulk domain reactions. Each reaction law is supplied with a chemical equation and a set of parameters and ChemChaste calculates the reaction rate, *R*(**u**).

**Table S1:** Reaction rates currently implemented in ChemChaste. The rates include constants *k_f_*, *k_r_*, *k_cat_*, *K_M_* and labels for spectator chemicals.For the reversible reactions, the rates may be negative 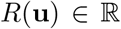 implying the reaction occurs in the reverse direction while irreversible reaction the rate must be positive 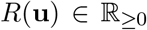 The user may implement their own reactions rates building upon these forms by adding a new reaction file to the inheritance structure of ChemChaste (Section S3).

| Reaction name | Reversible | Reaction rate formula $R(\mathbf{u})$ |
| --- | --- | --- |
| **ZerothOrderReaction** | False | $k f$ |
| **ZerothOrderReversibleReaction** | True | $k f - k r$ |
| **MassActionReaction** | True | $k f \prod_{c \in P} u_c^{\alpha_c} - k r \prod_{c \in S} u_c^{\beta_c}$ |
| **SpectatorDependentReaction** | False | $k f \prod_{c \in \tilde{S}} u_c$ |
| **MichaelisMentenReaction** | False | $k_{cat} u_E \prod_{c \in S} u_c^{\beta_c} / (K_M + \prod_{c \in S} u_c^{\beta_c})$ |

We use an inheritance strategy to readily build more complex reactions into ChemChaste, Figure S2. Therefore the functionality of the base reaction, `ZerothOrderReaction`, is inherited by all the upstream classes. The reactions currently implemented are considered foundational as they focus on different reaction properties (i.e reversibility, rate dependent on spectator species etc.) and may be easily built upon by a user to combine these properties under different rate laws.

**Figure S2:**
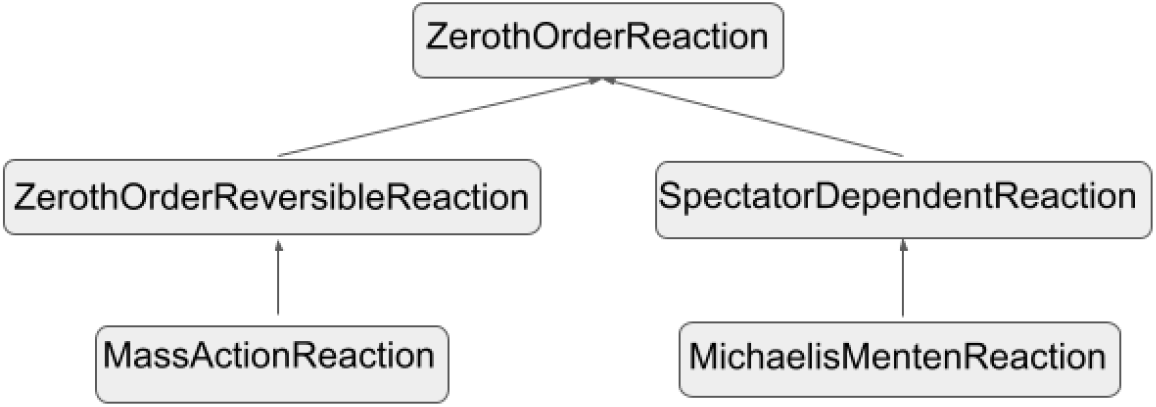
The inheritance structure for the bulk domain reaction files currently implemented in ChemChaste. These reactions are shown in Table S1. The structure builds from the base reactions, `ZerothOrderReaction`, to add more complex reaction rate laws.

#### S1.4 Simulations of coupled cell-domain systems: transport, *T*(u, u′), and membrane reactions, *M*(u, u′)

Simulations that couple a cell mesh to the reaction-diffusion domain utilise three different reaction types; chemical reactions (Table S1), transport reactions (Table S2), and membrane reactions (Table S3). The necessary file structure to call these reaction systems is given in Figure S9a. The transport reactions model chemical transport across the membrane and directly couple the cells to the domain, see Figure S1. Let the set of transported chemicals be denoted by *C* and a single chemical by *c* where *c* ∈ *C*. Let *C_domain_* denote the set of chemicals in the transport process located in the domain external to the cell membrane and *C_cell_* be the set of cellular chemicals. As the point *x* ∈ Ω is associated with both the domain and the cell there are two concentration vectors tied to the point, **u** for the chemical concentrations in the domain and **u′** for the concentrations within the cell. The lengths of these two vectors need not be the same, as in general |*C_domain_*| ≠ |*C_cell_*|.

Let the rate of the transport process connecting the two concentration vectors be denoted by *T*(**u, u′**)

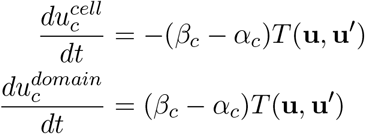

where 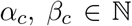 are the stoichiometric coefficients for chemical *c* ∈ *C* when considering the transport process as a “reaction”. Here *α* is the set of coefficients for the chemicals on the domain side of the membrane, that is the quantity of each species consumed in the forward sense of the process (i.e cell uptake), and *β* is the set of coefficients for the chemicals on the cell side of the process, (i.e cell excretion). For example, consider the reaction occurring at a rate *T*(**u, u′**);

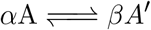

where *α* denotes the amount of *A* in the domain that are consumed in the forward reaction to produce *β* of *A*′ within the cell.

**Table S2:** The foundational transport process types implemented at present, including whether a process is reversible and the functional rate law utilised. The user may implement their own laws by adding a new transport reaction file to the ChemChaste system (Section S4).

| Transport process name | Reversible | Transport rate $T(\mathbf{u}, \mathbf{u}')$ |
| --- | --- | --- |
| ZerothOrderTransportIntoCell | False | $kf$ |
| ZerothOrderTransportOutOfCell | False | $kr$ |
| ZerothOrderReversibleTransport | True | $kf - kr$ |
| MassActionTransportReaction | True | $kf \prod_{c \in C_{cell}} u_c^{\alpha_c} - kr \prod_{c \in C_{domain}} u'_c{}^{\beta_c}$ |

The rate constants are defined with appropriate units, such that the units for the transport process are given in ’concentration per unit area per unit time’, i.e the amount of substance passing through the membrane in an infinitesimal length of time.

For reactions defined at the membrane, two separate reactions occurring on either side of the cell membrane, i.e. inside and outside, are coupled. Such reactions do not result in any transfer of chemicals across the cell boundary, but they alter the concentrations of species in the domain and inside the cell. The membrane reactions use two sets of stoichiometic coefficients. Let (*α, β*) be the stoichiometric coefficients for the reaction internal to the cell and (*α′, β′*) be the coefficients for the external domain reaction. As before, we denote the membrane reaction rate denoted by *M*(**u, u′**);

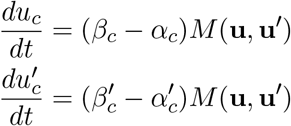

where variables and parameters take on the meaning defined previously. For two general bi-molecular reaction systems coupled at the membrane, the system takes the form;

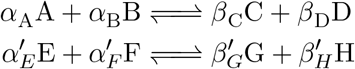

where the first and second reaction occur in the domain and inside the cell, respectively. The concentrations are provided by the separate state vectors as in Section S1.3 but with a shared rate, *M*(**u, u′**), which depends on both state vectors.

**Table S3:**
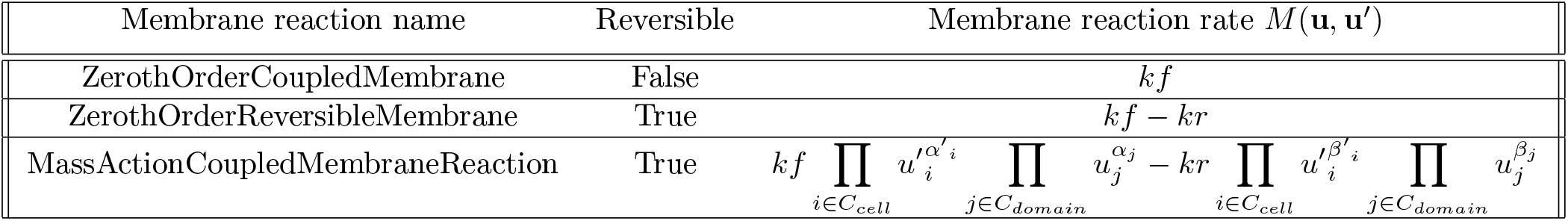
The membrane reaction types implemented at present, including whether a reaction is reversible and the functional rate law utilised. The user may implement their own membrane reaction laws by adding a new reaction file to the ChemChaste system (Section S5).

**Figure S3:**
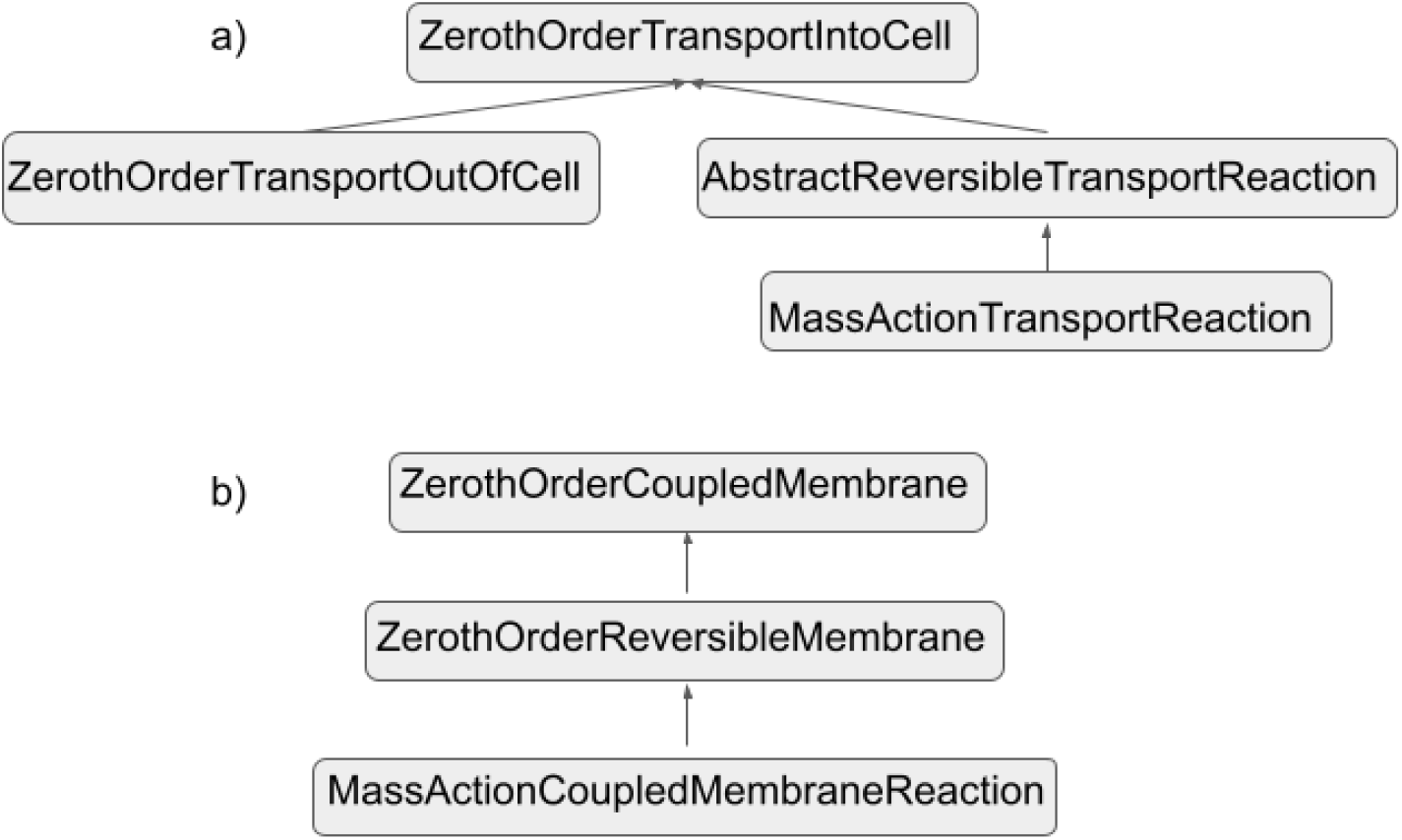
The inheritance structure for the transport a) and membrane b) reaction types currently implemented in ChemChaste. These reactions are shown in Table S2 and Table S3 respectively. a) The transport reactions build upon `ZerothOrderTransportIntoCell` as a base while in b) the membrane reactions build upon `ZerothOrderCoupledMembrane`. These laws may be built upon to add more complex transport and membrane reaction rates. The user may write their own transport processes and membrane rate laws by following the file structure given in Sections S4–S5.

### S2 Setting up a ChemChaste simulation with user-defined model properties and parameters

ChemChaste has been developed with the user in mind and simulations are defined in three steps. (i) The overall simulation parameters (Table S4) and file paths are specified in a configuration file; example configuration files are given in Figures S5 and S8. (ii) The reaction-diffusion system in the bulk is defined. The name of the directory is specified within the previous simulation configuration file (see Figure S5b). The reaction system (Table S1) and bulk domain information are stored in TXT and CSV files; exemplars are given in Figures S7 and S6 respectively. (iii) If coupling to a discrete cell-based model, the cell population and individual cell type properties are specified. The cell model information is given a separate directory within the simulation directory, as shown in Figure S9 for a two cell type example. For each cell type given in the TXT files, a cell-based reaction system is defined (Table S1) with cell-specific transport laws (Table S2) and membrane bound reactions (Table S3); example cell reaction files are given in Figure S10. The topology of the cell layer, i.e. the initial location of cells at the start of a simulation, is supplied by a CSV file together with a key file translating the cell ID used in the topology file to the names of the cell types (which are also used in the file directories for cell files).

In the following sections, we explain each of the required configuration files and their contents to run a ChemChaste simulation. These configuration files are parsed and fed into a C++ ChemChaste simulation through the help of a “run-script”.

#### S2.1 ChemChaste run script

To run a ChemChaste simulation, users can make use of a “run-script” file, which is a command line script written in Phyton 3 language (see example in Figure S4). A new “run-script” file should be supplied for each simulation, although simulations with same files but different parameters, e.g. for parameter ’sweeping’, can be provided through the same “run-script” file (see example). The run-script sets the ChemChaste executable to run in naive parallel, that is one simulation per processor, and runs the testing and compilation features of Chaste. Within the “run-script” file the user defines the relative directories for the different configuration files, as well as some of the global simulation parameters. The full set of parameters that can be set in the “run-script” file are given S4. Here, the configuration files for the Cross feeding (Section S2.3) and Schnackenberg spatial patterning (Section S2.2) are used as examples. In the example file provided, a total of 10 simulations are executed, 5 of type `complex_cell` and 5 of type `domain_only`, and distributed over 3 cores. Some of these parameters can also be set in other configuration files, as discussed below.

**Figure S4:**
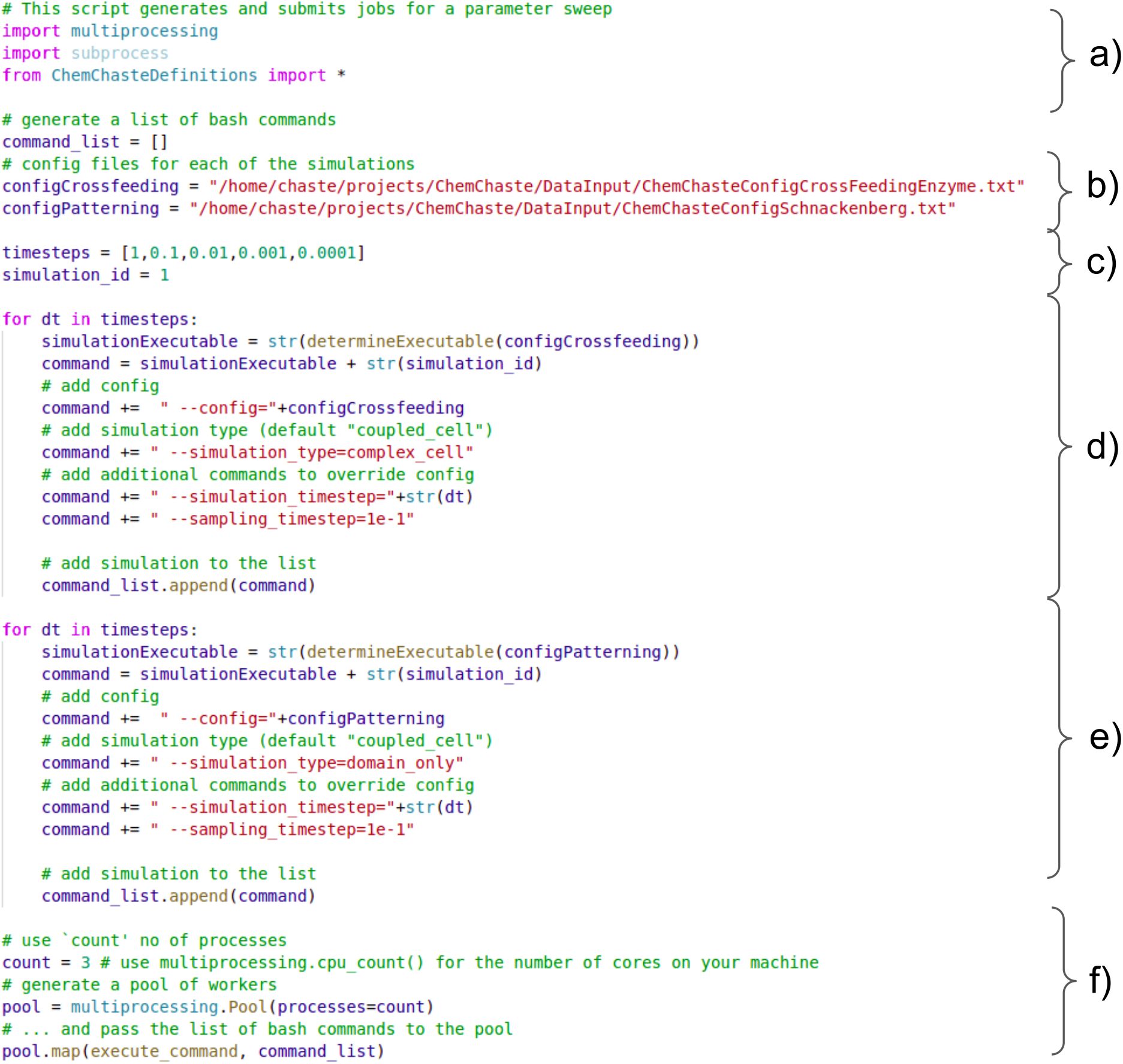
Example “run-script” file for defining features of ChemChaste simulations. The structure of this “run-script” file is such that it is divided into sections, as shown with a letter-based labelling on the figure and as explained next. a) This section defines the import files used for running parallel simulations and controlling the simulation compilation and implementation. b) This section defines the configuration files for each user simulation that are contained within the *DataInput* directory of ChemChaste. c) This section defines the global simulation parameters, in particular those that are used for parameter sweeping. The simulation ID created in this section is shared across a set of simulations to be run in parallel. For these simulations, a different parameter value is to be used - in this example the “time step size” parameter. d) This section defines the command to be used in the command-line initiation of a simulation. The command is created in a series of steps, by appending different aspects of the simulation command together. First the simulation executable, in this case the cross feeding simulation, is created and then appended the simulation type. Then, the desired simulation parameters are amended to the command. In this case, note that the simulation timestep is set by grabbing its value from the provided parameter list and by making use of a for loop structure. Finally the destination for the data file is appended to the command using the parameter `sampling_rate` and a value of 1*e*^−1^. e) This section is a repeat of section d) but using a reaction only simulation example. The Schnackenberg simulation is used with the same parameters as in d) but where `simulation_type` is set to `domain_only`. f) This section defines final aspects of simulations, such as number of processor cores. The parallel simulations are mapped to `count` number of processor cores.

**Table S4:**
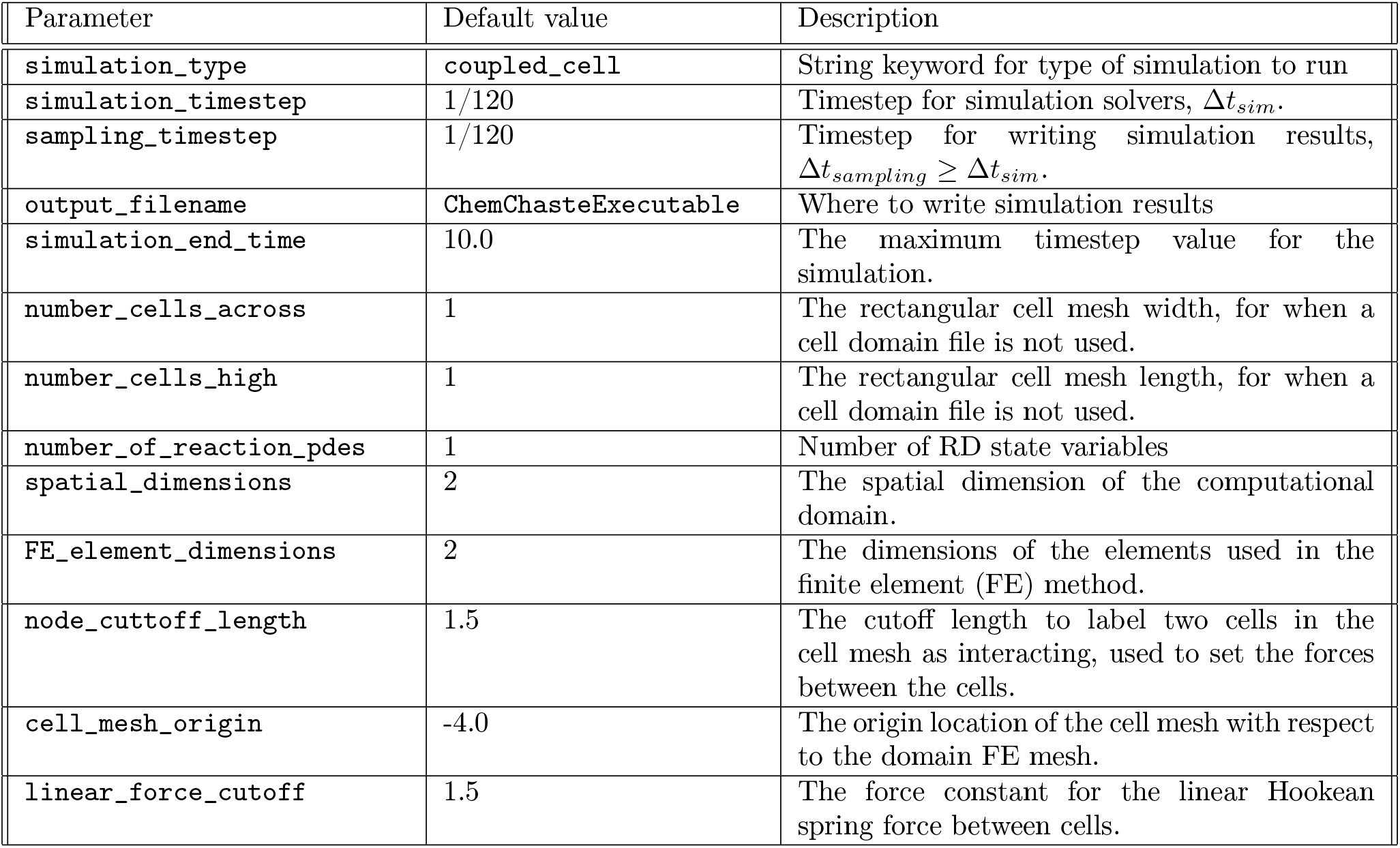
ChemChaste simulation parameters that can be set via the “run-script” file. These parameters control the type of simulation to run, the solver properties such as end time and solver time step, and finite element properties such as spatial domain dimensions and element dimensions for the FE implementation. Each parameter is set by name in the simulation configuration file, an example is provided in Figure S5a.

Among the different parameters that can be set in the “run-script” file, and listed in S4, we highlight here some of the key ones. The `simulation_type` parameter, which sets the solver methods used in ChemChaste. This parameter may be specified in either the configuration file or in the run-script (Figures S5 and S4). The parameter options; `domain_only`, `coupled_cell`, `complex_cell`, control how the ChemChaste system builds the simulations. The `domain_only` option is used to solve a reaction-diffusion system in a domain without cells (removing the summation term from equation (1)), while the coupled cell-domain simulations are simulated using either the `coupled_cell` model and the `complex_cell` model. These two models share the same parameter sets and file systems, see Table S4 and Figure S9, but differ in the cell division implementation. The `coupled_cell` simulations model the cells divide into a parent and offspring cell. The concentration contents of the parent cell are duplicated and copied over to the offspring cell. In contrast, for the `complex_cell` model the concentration contents of the parent cell are either divided equally between the parent and offspring or duplicated and a further “speciesDivisionRules.csv” file is needed for each cell type to control the sharing behaviour.

The cell population may be defined using the file system (see Section S2.3) or by providing the population size. The size is provided through the `number_of_cells_high` and `number_of_cells_across` configuration parameters. These parameters are used to construct a honeycomb mesh of length `number_of_cells_high` and width `number_of_cells_across` with the origin of the mesh compared to the PDE domain provided by `cell_mesh_origin`. The `cell_mesh_origin` value is added to the *x* and *y* direction to translate the positions of the nodes in the cell mesh. The edges of the mesh denote which cells are nearest neighbours and whether these cells interact depends on the distance between the cells. These neighbours interact if their positions are within the cut off distance, `node_cuttoff_length`, and the strength of the linear Hookean force for the interaction is provided by `linear_force_cutoff`.

#### S2.2 Configuration files for Domain only simulation

For reaction-diffusion simulations, the user provides a set of files specifying the domain topology, boundary conditions (BCs), initial conditions, and reactions systems (see examples in Figures S6–S7). These files are provided to ChemChaste by defining their file paths in the configuration file and an associated directory structure (see examples in Figure S5a Figure S5b).

**Figure S5:**
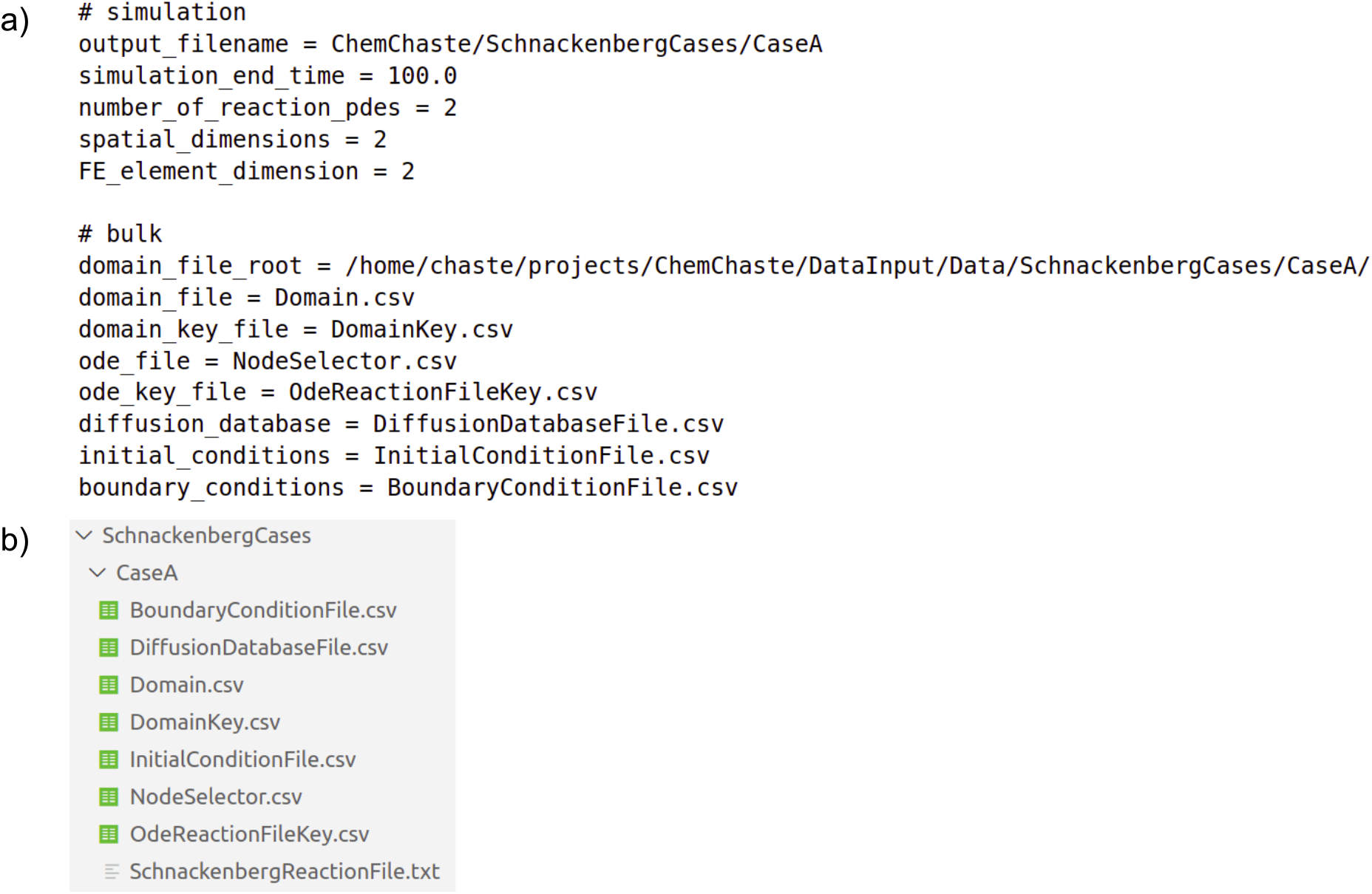
The configuration file a) and overall directory structure b) for simulating a reaction-diffusion system. a) The configuration file containing the basic simulation parameters; output directory, simulation end time, number of chemical PDEs to simulate, the domain and FE element dimensions. The configuration file also contains the directory paths and names of the different files used in the simulation. The file structure used during the domain only simulation b). The file paths are defined within the configuration file and follow CSV file types for parameters and defining the domain while TXT files are used for writing the reactions.

##### S2.2.1 Building the domain with chemicals and setting the properties

Users can define the domain in a series of CSV files.

“Domain Key” and “Domain Information” CSV files: These two files together define the domain topology. The “Domain Key” file introduces numeric id’s for different ’types’ of sub-domains, which might be associated with different chemical diffusion rates. Here, we use an example to define two sub-domains within the domain as ’bulk’ and ’film’, e.g. to mimic a bulk and biofilm environment. The “Domain Key” file simply lists names for such sub-domains and associates them with a numeric id. In the example given in Figure S6a-c, we defined two sub-domains labelled as ’Bulk’, id 1, and ’Film’, id 2, and then distributed them on the domain in such a way that the left section is ’bulk’ and right section is ’film’. Note that the “Domain Information” file is organised as a 2D matrix, which is mapped on to the mesh implemented in the FE simulations. This mapping stretches each matrix entry into 10 elements in the FE mesh. That is, a 10×10 file matrix would map onto a 100×100 FE mesh.

“Boundary Condition” and “Diffusion Database” csv files: Chemicals that diffuse and react within the domain are first defined in a “Boundary Condition” file. This file introduces each chemical in the system and sets the boundary conditions for them. In the example given in (Figure S6, we define two chemicals, *U* and *V*, and set the boundary conditions for both as zero-Neumann (zero-flux). Each chemical’s diffusion in the different sub-domains (in this example, ’film’ and ’bulk’) are then defined in a file called, “Diffusion Database”. Here, the user can use the defined sub-domain names (see above) to then specify the diffusion rate for each chemical in that sub-domain (see Figure S6d). Occasionally, a model may require the diffusion of a chemical to be completely inhibited in a specific sub-domain, this may be done by setting the diffusion rate value to 0.

“Initial Condition” csv file: The initial concentrations of each chemical in each sub-domain are given in a file called ’Initial Condition’. The file should specify one initial concentration value for each chemical and for each sub-domain (Figure S6e). This initial value is then applied to all mesh nodes associated with that sub-domain (see equation 4. While this approach provides a homogenous condition across all nodes of a given sub-domain, the values at the individual nodes may be perturbed through the addition of random noise value defined on a nodal basis during the setup of the simulation. That is, for states *U, V* with initial value *u*_0_, *v*_0_ for a given sub-domain, Ω*_sub_* the perturbed initial values *U*(*x*, 0), *V*(*x*, 0) are given by;

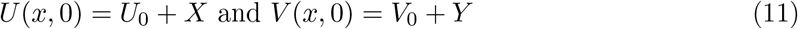

where the node location *x* ∈ Ω*_sub_* and *X, Y* ~ *Uniform*(−1, 1) are uniformly distributed random noises on the interval [−1, 1]. This random perturbation occurs only if the perturbation option in the initial condition file is set to `true` (see Figure S6e).

**Figure S6:**
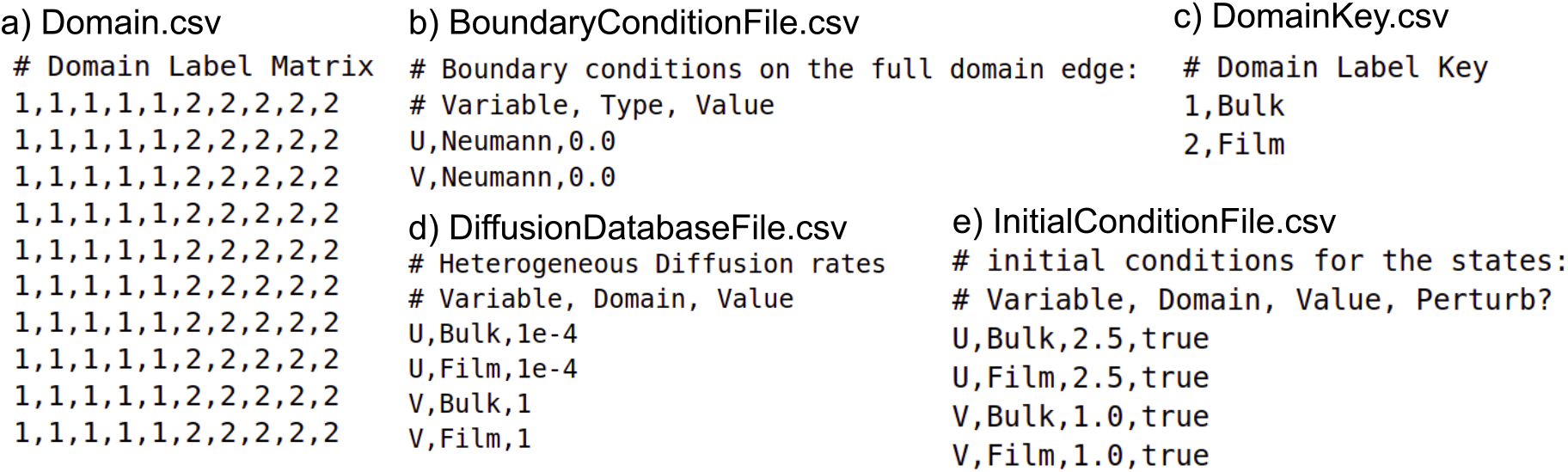
Files defining the domain topology and providing simulation parameters. a) CSV file defining the domain topology. The nodes on the domain are labelled according to this matrix where sub-domains are specified by using different labels. The size of the matrix specified in this file is directly proportional to the size of the simulation mesh after scaling. Therefore a rectangular domain specified in this file will produce a rectangular domain mesh. b) Boundary conditions are implemented on the outside nodes of the FE mesh. The conditions are specified as “state variable”, “BC type”, and “BC value”. c) Domain key file, which connects the ‘simple’ labels used in the domain topology file with the domain names. d) Diffusion in the simulations is modelled as isotropic diffusion where the coefficient value is the same in all directions. These values for each state variable are provided for all sub-domains names in c). e) The initial conditions in the domain are defined for each state variable (chemical) on each sub-domain. The value can differ for each sub-domain and the user can specify whether to perturb the value by a uniform random value.

##### S2.2.2 Chemical reactions and the “Reaction Information” and “ODE Reaction Key” CSV files

To define domain reactions the user provides a set of files, similar to those used for defining sub-domains. The user first creates a file called “ODE Reaction Key”, specifying a reaction file name and an associated numeric id for that reaction (see Figure S7b). The reaction id can then be used to specify the nodes that are associated with specific reactions in the domain. This information is provided in a file called “Reaction Information”, which should provide a matrix of the equal size to that given in the “Domain Information” file, where the reaction id’s on the nodes are listed (Figure S7a). In the provided example, we have defined a reaction with reaction id 0 and attached this reaction to all of the nodes in the domain. Details of each of the implemented reactions are provided in a separate TXT file, which is referred to in the “ODE Reaction Key” file. In this example, we called this file “Schnackenberg Reaction” file. This file describes the reactions among the chemical species (see previous section), as well as the associated reaction parameters (Figure S7c). This description file utilises the provided reaction types and reaction parameters, as detailed in Section S1.3. Note that, as seen in the provided example, the file describing the ODE reactions can feature multiple reactions and all of the reactions defined in the file will be performed on the associated nodes (see Figure S7).

**Figure S7:**
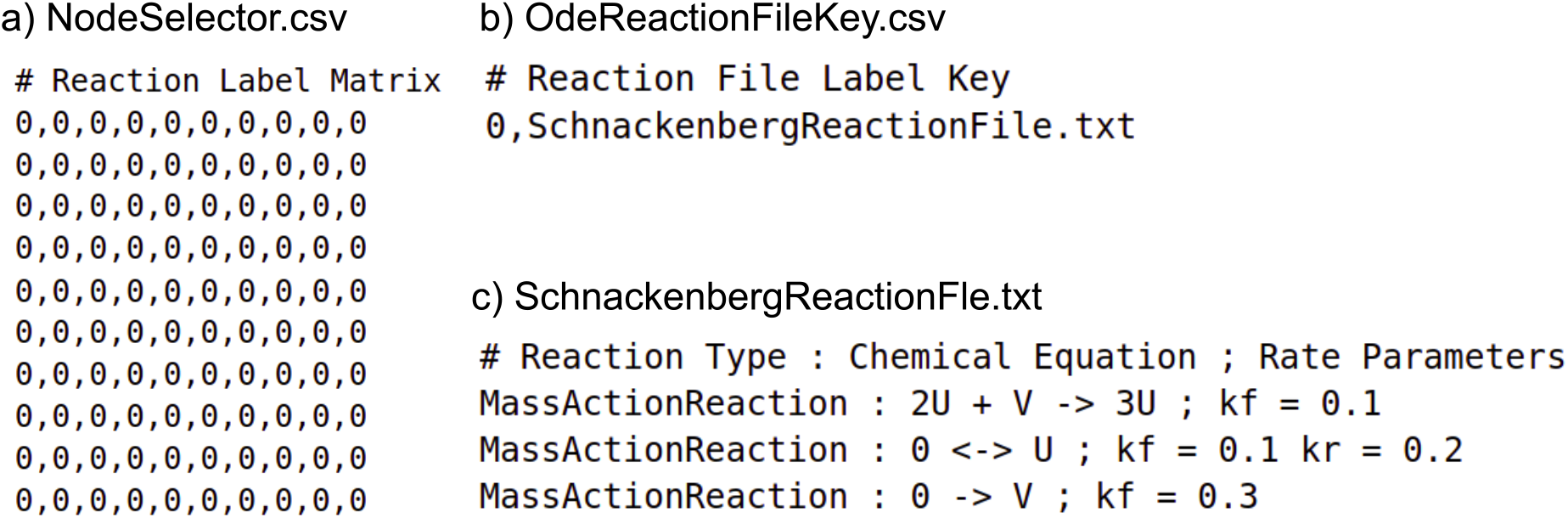
The implementation of bulk reactions in user defined files. a) A CSV file defining the numeric ID of reaction files, which describe a series of chemical reactions. The reactions defined in a file associated with a node will determine those reactions to be active on that node. This allows for the creation of reaction sub-domains, which are not necessarily the same as the diffusion sub-domains. b) A reaction key file that connects the numeric IDs used in part a) to actual reaction file names. These names refer to TXT files containing the reaction system that are to occur on the associated nodes (creating the reaction sub-domain). c) An example reaction file, `SchnackenbergReactionFile.txt`. Each line denotes a separate reaction. Reactions are defined using a standard form composed of: “rate law”, “ : ” rate delimiter, reaction equation, “ ; ” reaction delimiter, and the rate law parameters.

#### S2.3 Defining a cell simulation - Cooperator-cheater system

The hybrid continuum-discrete models of cells coupled to a bulk require a modified file structure and additional files. In particular, the cell-coupled simulation defines an additional cell mesh whose nodes denote the cell locations. The cell nodes are associated with a cell type and a cell object is formed with the cell properties and reactions corresponding to that cell type label. The simulation parameters for a cell-coupled simulation are presented in Figure S8 with the directory file structure given in Figure S9. The directory structure is such that the domain files are stored within a `DomainField` sub-directory of the simulation directory (Figure S9a).

**Figure S8:**
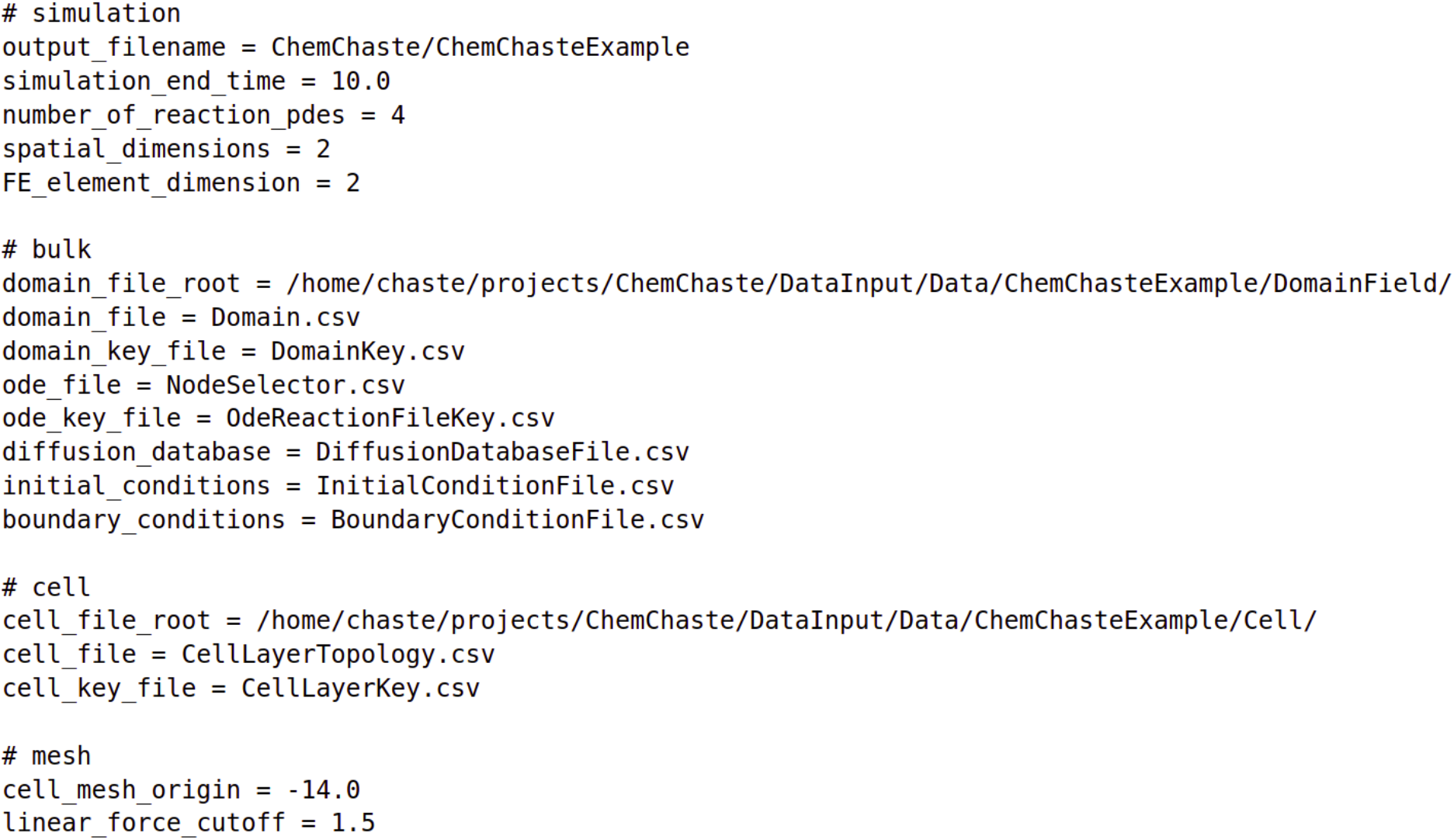
The configuration file for the cell-coupled simulations. This configuration file defines the directory paths to associated files and includes some parameters specific to cell-coupled simulation. The files, accessible via the defined directory paths, define the structure and sub-populations of cells; the information is stored in the files `cell_file_root`, `cell_file`, `cell_key_file`. See example given in Figure S10. Simulation parameters used to couple the cell and domain mesh are also provided. “`cell_mesh_origin`” denotes the origin of the cell population structure with respect to the domain mesh. “`linear_force_cutoff`” is used as an interaction strength parameter for the Hookean linear spring force which connects the cells in the simulation.

##### S2.3.1 Including cells into the simulation

The `coupled_cell` and `complex_cell` simulation types add cells to the reaction-diffusion domain. Each cell has a type and the properties of each type are stored within a series of files contained in a directory, Figure S9a. The population is given a structure by constructing a mesh and associating a cell object to each node. Example files defining the mesh dimensions and cell-node labelling can be seen in Figure S10a–b. The initial structure of the cell population, at simulation start, is provided as a matrix in the `“CellLayerTopology”` CSV file (Figure S10a). The matrix is required to be smaller than that of the domain (Figure S6). The entries in this matrix refer to the type of a single cell which are mapped onto honeycomb mesh of equal length and width to the input matrix. If an input matrix is not specified and instead the configuration parameters, `number_cells_high` and `number_cells_across`, are provided then these values are used for the length and width of the honeycomb mesh. The cells are labelled by a numeric ID with cell type names provided in the `“CellLayerKey”` file. In the example shown in (Figure S10 two cells are defined with one cell of each type in the domain and are then provided the names ’CellA’ and ’CellB’. These names are also used as sub-directory names for the following cell specific files. The initial cell concentrations and the concentration thresholds for the chemicals within the cell are given with a separate file for each cell type (see Figure S9b–c). The concentration thresholds provide a lower and upper bound for each of the chemical concentrations, which are used to implement cellular ’rules’. If the cellular concentration falls below the lower threshold the cell is marked for death while cells with concentrations above the upper threshold will undergo cell division. The cell reaction systems are provided (Sections S1.3 and S1.4) see Figure S10c–e. The reactions occurring within the cell are located in the “Srn” text file while the transport and membrane reactions are located in the “TransportReactions” and “MembraneReactions” files.

As with the nodal initial conditions for the reaction-diffusion simulation, the initial conditions for the cellular concentrations may be perturbed by adding a random noise value at the beginning of the simulation. For the concentration *u*(*t*), with initial concentration *u*_0_ and with the perturbation option set to `true`, the starting cell concentration is given by;

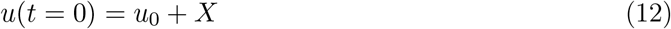

where *X* ~ *Uniform*(−1, 1) is uniformly distributed random noise on the interval [−1, 1].

During the simulation the conditions for cell division and death are determined by the species threshold file, Figure S10c. Each chemical in the cell reactions is provided with a maximum, *u_max_*, and a minimum, *u_min_*, concentration threshold. If the concentration reaches the threshold *u_max_* for at least one chemical the cell division process is triggered. If the simulation type is set to `coupled_cell` then the cell concentrations are duplicated and copied during division. For the `complex_cell` simulation the cell contents of the parent are shared with the offspring. That is for each cell chemical *c* of parent concentration *u_c_* at division the new concentrations 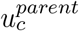 and 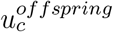 are given by

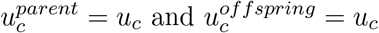

for the `coupled_cell` simulation type and for the `complex_cell` simulation type

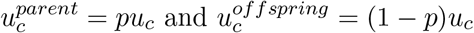

where *p* is the splitting ratio (*p* = 0.5 by default). Cell death is implemented by the removal of a cell and its associated cell mesh node. This apoptosis process is triggered when the concentration of a chemical falls below the minimum threshold for that species, *u* ≤ *u_min_*. To remove this death functionality, users should set *u_min_* = 0.0. This will prevent the apoptosis process being triggered by that chemical. To remove the cell division functionality, users should set *u_max_ ≤ u_min_* or *u_max_* = 0.0.

In the example presented in Figure S9 we define two cell types, {*CellA, CellB*}, and the corresponding cell directories. The initial conditions and species thresholds are presented for *CellA* covering each chemical found in the reactions, Figure S9. The cell division/death processes are dependent on the threshold values for each chemical but this has been set to ignore all of the chemicals, by setting both the upper and lower thresholds to zero, except for `Biomass` which will trigger division at a concentration *u_max_* = 1.5 and apoptosis at *u_min_* = 0.1.

**Figure S9:**
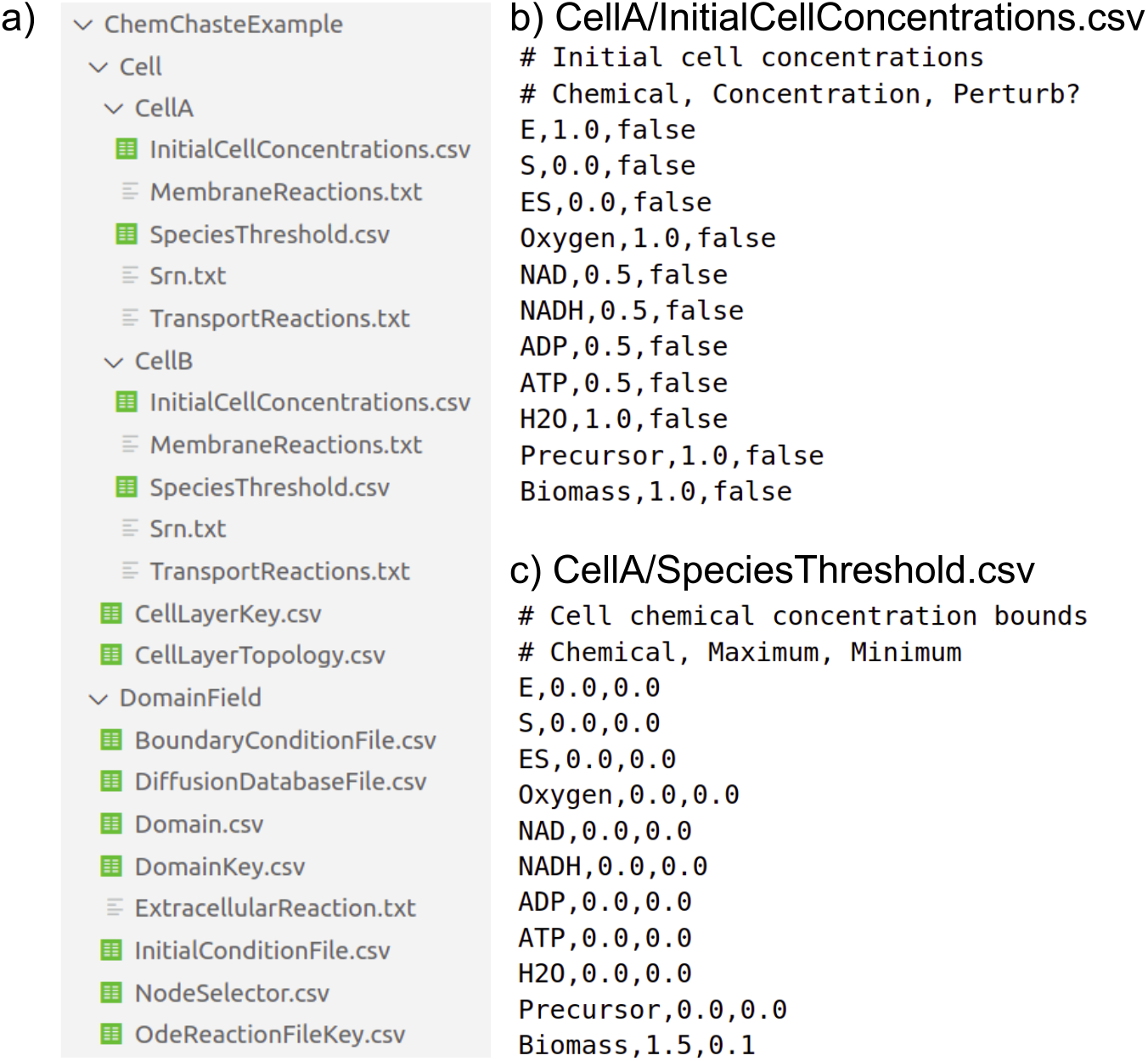
Description of directories and cell properties files for the cell-coupled simulations. a) The directory structure for a cell-coupled simulation. Files relating to the domain structure, as detailed in Figure S6, are contained within `DomainField` directory. The cell files are provided within the `Cells` directory. Each cell type is provided its own sub-directory, with the same name as the label name in the `CellLayerKey.csv`, Figure S10d. For each cell type we define an initial concentrations file b) and a species threshold file c). b) The initial conditions are provided for each cellular state variable (chemical) following the form; name, value, and whether to perturb the initial value on a nodal basis. c) The threshold values for each cellular state variable (chemical) are provided in the order; name, maximum value, minimum value.

**Figure S10:**
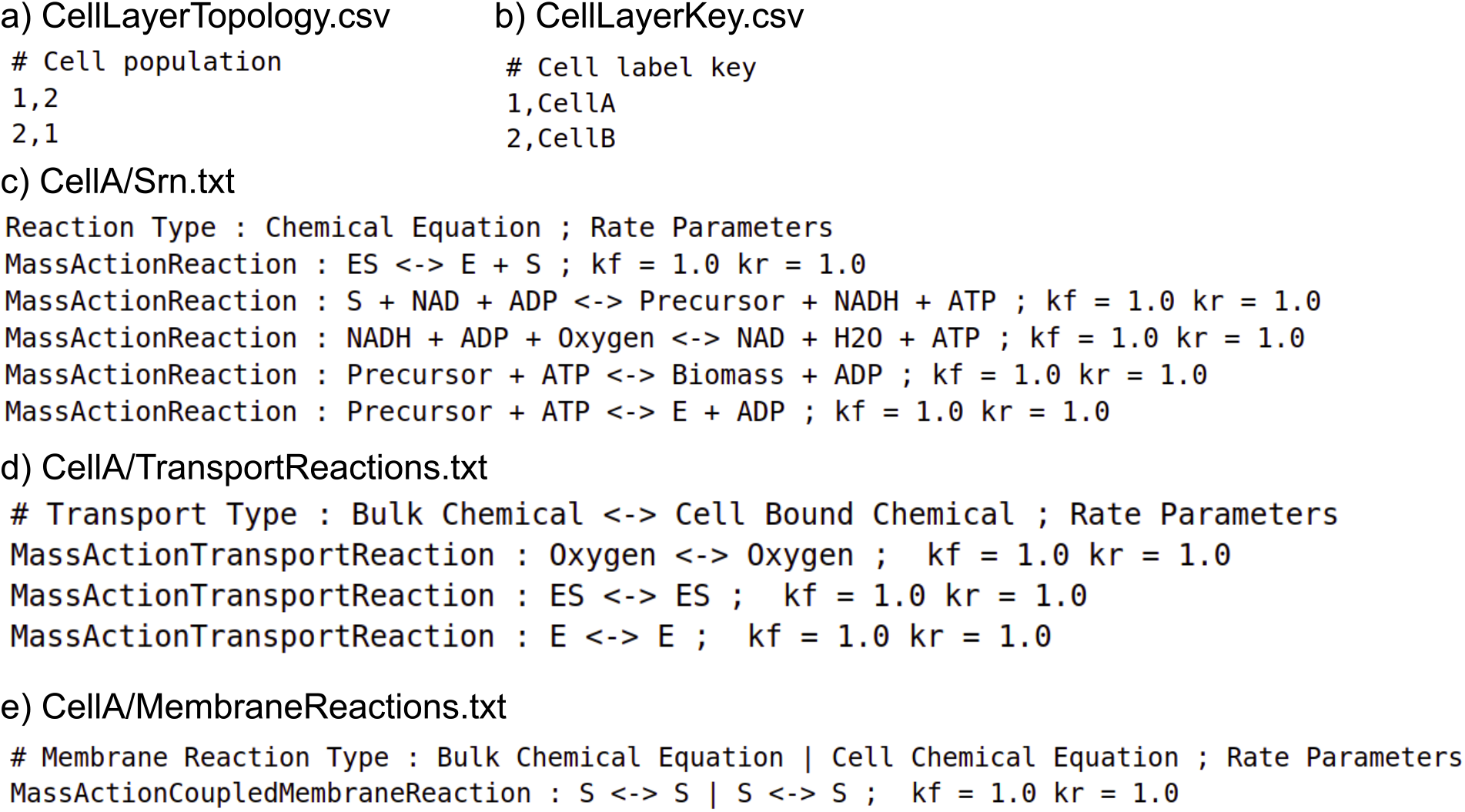
The files associated needed to form the cell mesh and populate the cells with chemical reactions. These files are to be placed in the Cell sub-directory of the ChemChaste main directory (see Figure S9a). a) The `“CellLayerTopology.csv”` file provides the information for the initial cell mesh topology. It is written in the same format as the domain layer FigureS6a. In this example, a rectangular mesh of two cells is defined where the first cell is labelled `“1”` and the second labelled `“2”`. This mesh is aligned with the domain mesh through translating the origin of the cell mesh as specified by the `cell_mesh_origin` parameter in the configuration file (see Figure S8). b) The `“CellLayerKey.csv”` files contains the key mappings from the numeric ID label used in `“CellLayerTopoogy.csv”` to the cell type used for the directory names. In this example two cell types are used *CellA, CellB*. The cellular reactions for *CellA* are provided in three TXT files; c) the internal reactions, d) the transport reactions, and e) the membrane reactions. c) The `“Srn.txt”` file contains the cellular reaction system. Each line contains one reaction. The reactions are written in the standard form; reaction kinetic law, “ : ” reaction law delimiter, reaction chemical equation, “ ; ” parameter delimiter, kinetic law parameters. d) The file `“TransportReactions.txt”` lists the reactions/processes that couple the cells to the external domain. The reactions in this file are written in such a way that the chemicals on the left hand side represent the species in the bulk domain, while those on the right hand side represent the species inside the cell. Otherwise they follow the form in c) but using an appropriate set of reaction rate laws. e) The file `“MembraneReactions.txt”` lists the reactions that are coupled at the cell membrane and with reaction occurring on outside of the cell and one on the inside of the cell. These reactions are separated by the membrane delimiter, “ | ”, when written and the reaction rate law belongs to the membrane reaction set.

### S3 Adding a new reaction rate law

Chemical reactions occurring in the cell or in the domain follow a set of reaction rate laws defined in Table S1. However, new chemical reaction types may be added by the user and then freely implemented within the reaction system files. To introduce a new reaction rate, a new reaction header, which inherits from a previous reaction class, needs to be created and placed into the inheritance hierarchy (Figure S2). The user would then populate the class, update the inherited virtual functions, and then update the `ReactionTablet` function within the `ReactionTypeDatabase` file. From there on, the reaction rate type may be called by name, in the same manner as with the in-built reaction rates; e.g `MassActionReaction`.

The reaction header file needs to include the falling classes as “includes”; `AbstractChemical`, `AbstractChemistry`, and `AbstractReaction`. This will allow the class definitions to inherit publicly from the core reaction types. For example, `AbstractReaction` for a general irreversible reaction or `AbstractReversibleReaction` for a general reversible reaction. This inclusion is needed (useful), since these inherited reaction types provide a set of virtual functions that are useful to construct the reaction mechanism. In the following, we briefly describe these virtual functions, which the user will have to consider changing when creating a new rate law.

1. `React()`
  - Function description: Virtual function that implements the core dynamics of a chemical reaction. This function takes in the current system concentrations and outputs a vector describing the change in concentrations for the next timestep. The function calls `UpdateReactionRate()` function and then applies the reaction rate to the reaction stoichiometry, so to calculate the change in the species concentration.
  - Function input variables:
    – |systemChemistry (`AbstractChemistry`*)|
    – |currentChemistryConc (const std::vector<double>&)|
    – |changeChemistryConc (std::vector<double>&)|
  - Note: It might not be necessary to change this virtual function when creating a new rate law. This is because calculating the change in the chemical concentrations as the product of the reaction rate and stoichiometry vector is a standard method for implementing chemical reactions in a dynamical simulation. However, this function is explained here so that users are aware of it and can have the flexibility to implement other modelling approaches to chemical reactions by changing it.
2. `UpdateReactionRate()`
  - Function description: This function calculates the reaction rate scalar value for a given reaction. If the reaction being operated on is an irreversible reaction inheriting from `AbstractReaction`, then this function calculates the forward reaction rate, `forward_rate`, and calls the function `SetReactionRate(forward_rate)`. If the reaction being operated on inherits from `AbstractReversibleReaction` and has a calculated reverse reaction rate, `reverse_rate`, then this function calls both of the functions `SetForwardReactionRate(forward_rate)` and `SetReverseReactionRate(reverse_rate)`.
  - Function input variables:
    – |systemChemistry (AbstractChemistry*)|
    – |currentChemistryConc (const std::vector<double>&)|
  - Note: This function is the main function to modify when creating a new reaction rate type. It essentially determines how reaction rates are calculated from current system concentrations and stoichiometries. The `MassActionReaction` class, which is of the reversible reaction type, utilises the current system concentrations to calculate reaction flux values in both reaction directions. The reaction rate values are calculated for use in the `React()` method (see Table S1.
3. `GetReactionType()`
  - Function description: This function returns a string type which provides the name of the reaction type; for example returning `MassActionReaction` or `SpectatorDependentReaction`. This function may also be used for reaction tracking purposes, but in the current implementation, it is used in the `ReactionTablet` function. This name needs to be the name in which the reaction files label the reaction type in order for the correct class calls to be made.
4. `UpdateReaction()`
  - Function description: This is a void function with no inputs. It is provided for the case that a reaction’s behaviour needs to be altered outside of the `React()` function. Possible utilities include a switching of behaviour in reaction style based on concentrations, time, or system properties.
5. `ParseReactionInformation()`
  - Function description: This function parses the parameters and variables needed to process a reaction. In terms of the written file reaction, these data values occur in the string after the ; delimiter. This string is parsed into the data values provided by the user using a string delimiter. The user needs to identify the delimiter of this string as a member value in the reaction class. The values parsed are to be also stored as member values and may be utilised in the `UpdateReactionRate()` function where necessary.
  - Function input variables:
    – `reaction_information (string)`
    – `IsReversible (bool)`

### S4 Adding a new transport process law

The cells and the reaction-diffusion domain are coupled through the transport processes transferring chemical species to either side of the cell membrane, see Section S1.4. As these processes require the chemical concentrations of both the cellular species and the corresponding external species at that domain location to be known, transport rules have a different construction to the reaction rate laws described in the previous section. ChemChaste has a set of transport rates already defined, Table S2, but new transport process rates may be added by the user and integrated with ChemChaste.

Transport processes in ChemChaste follow the general form of a reaction where the `Substrates` are chemical species in the domain and the `Products` are species in the cell. For the introduction of a new transport process, users would need to create a new transport reaction header, which inherits from a previous transport reaction class (see Figure S3a for the inheritance structure for transport reactions).

Within the new transport reaction the user would populate the class with updated inherited virtual functions, then update the `TransportTablet` function within the `ReactionTypeDatabase` file. From there the reaction type may be called by name in the same manner as the in-built reactions; i.e `MassActionReaction`.

The reaction header file needs as ’includes’; `AbstractChemical`, `AbstractChemistry`, and the appropriate abstract transport reaction base. The available abstract bases are; `AbstractTransportReaction` for single direction domain to cell transport, `AbstractTransportOutReaction` for single direction cell to domain transport, or `AbstractReversibleTransportReaction` for reversible domain-cell transport. Depending on the abstract base selection, the class definitions needs to inherit publicly from the base reaction types. These inherited base reaction types provide a set of virtual functions which control the reaction mechanism. In the following, we briefly describe these virtual functions, which the user will have to consider changing when creating a new transport rate.

1. `React()`
  - Function description: Virtual function that performs the transport reaction. This function takes in the current bulk domain and cell concentrations and computes a vector describing the change in both concentration sets over the next timestep, i.e. the reaction rate. The function calls `UpdateReactionRate()` function, which multiples the reaction rate with the reaction stoichiometry to update the species concentrations.
  - Function input variables:
    – `bulkChemistry (AbstractChemistry*)`
    – `cellChemistry (AbstractChemistry*)`
    – `currentBulkConcentration (const std::vector<double>&)`
    – `currentCellConcentration (const std::vector<double>&)`
    – `changeBulkConc (std::vector<double>&)`
    – `changeCellConc (std::vector<double>&)`
  - Note: It might not be necessary to change this virtual function when creating a new transport rate. The input variables `bulkChemistry` and `cellChemistry` refer to the chemical species outside the cell in the domain and inside the cell respectively. The `bulkChemistry` species refer to the `Substrates` of the transport reaction and the `cellChemistry` refers to the `Products` of the reaction.
2. `UpdateReactionRate()`
  - Function description: This function calculates the reaction rate scalar value for the given reaction. If the reaction being operated on is an irreversible reaction inheriting from `AbstractTransportReaction` or `AbstractTransportOutReaction`, then this function calculates the forward reaction rate, `forward_rate`, and calls the function `SetReactionRate(forward_rate)`. If the reaction being operated on inherits from `AbstractReversibleTransportReaction` and has a calculated reverse reaction rate, `reverse_rate`, then this function calls both of the functions `SetForwardReactionRate(forward_rate)` and `SetReverseReactionRate(reverse_rate)`.
  - Function input variables:
    – bulkChemistry (AbstractChemistry*)
    – cellChemistry (AbstractChemistry*)
    – `currentBulkConc (const std::vector<double>&)`
    – `currentCellConc (const std::vector<double>&)`
  - Note: This function is the main area to modify for new reaction types.
3. `GetReactionType()`
  - Function description: This function returns a string type which provides the name of the reaction type; for example returning `MassActionTransportReaction`. This function may also be used for reaction tracking purposes, but is currently used in the `TransportTablet` function. This name needs to be the same name in which the reaction files label the reaction type in order for the correct class calls to be made.
4. `UpdateReaction()`
  - Function description: Void function with no inputs provided for the case that a reaction’s behaviour needs to alter outside of the `React()` function. Possible utilities include a switching of behaviour in transport style based on concentrations, time, or system properties.
5. `ParseReactionInformation()`
  - Function description: This function parses the parameters and variables needed to process the transport process. In terms of the written file reaction, these data values occur in the space after the ; delimiter. This string is to be parsed into the data values by the user using a string delimiter. The user needs to identify the delimiter of this string as a member value in this reaction class. The values parsed are to be also stored as member values and may be utilised in the `UpdateReactionRate()` function where necessary.
  - Function input variables:
    – `reaction_information (string)`
    – `IsReversible (bool)`

### S5 Adding a new membrane reaction rate law

Besides reactions outside and inside the cell, chemical reactions may also be modelled as if occuring at the cell membrane in a way that couples cell interior and external chemicals. These reactions essentially couple two reactions, one occuring in the domain and another in the cell. These membrane reactions are described in Section S1.4, with the currently implemented membrane reaction rates in Table S3.

New membrane reaction rates may be added by the user by creating a new reaction class inheriting from an existing membrane reaction class (Figure S3b), overriding the inherited virtual functions. To integrate with the ChemChaste system the `MembraneTablet` function within the `ReactionTypeDatabase` file is updated with the new membrane reaction. From there the reaction type may be used by name in the same manner as the supplied membrane reactions; i.e `MassActionCoupledMembraneReaction`. As the membrane reaction couples two separate reaction systems, two reactions are provided per instance. These separate chemical reactions within the membrane reaction are separated by the | delimiter, with the external domain reaction before the delimiter and the internal cell reaction after.

The membrane reaction header file needs as “includes”; `AbstractChemical`, `AbstractChemistry`, and either of the base membrane reaction types `AbstractMembraneReaction` for irreversible reactions or `AbstractReversibleMembraneReaction` for reversible reactions. Coupling of a reversible and irreversible reaction is not currently implemented. The class definitions then inherit publicly from the appropriate base membrane reaction type. These inherited types provide a set of virtual function which control the reaction mechanism. In the following, we briefly describe these virtual functions, which the user will have to consider changing when creating a new membrane-bound reaction rate.

1. `React()`
  - Function description: Virtual function that performs the membrane reaction. This function takes in the current bulk domain and cell concentrations, and computes a vector for the change in both concentration sets over the next timestep. The function then calls the `UpdateReactionRate()` function, which applies the reaction rate to the stoichiometry to calculate the change in concentrations for both the domain and cell chemicals.
  - Function input variables:
    – `bulkChemistry (AbstractChemistry*)`
    – `cellChemistry (AbstractChemistry*)`
    – `currentBulkConcentration (const std::vector<double>&)`
    – `currentCellConcentration (const std::vector<double>&)`
    – `changeBulkConc (std::vector<double>&)`
    – `changeCellConc (std::vector<double>&)`
  - Note: Calling and modifying this function directly may not be necessary to implement a new membrane reaction rate. The input variables `bulkChemistry` and `cellChemistry` refer to the chemical species outside the cell in the domain and inside the cell respectively. These separate chemistries are formed from both the substrates and products of the separate chemical reactions.
2. `UpdateReactionRate()`
  - Function description: This function calculates the combined reaction rate scalar value for both the reactions at the membrane. After calculating the reaction rate, `forward_rate`, the function calls the function `SetReactionRate(forward_rate)` for an irreversible reaction inheriting from `AbstractMembraneReaction`. If the reaction inherits from `AbstractReversibleMembraneReaction` and has a calculated reverse reaction rate, `reverse_rate`, then both the functions, `SetForwardReactionRate(forward_rate)` and `SetReverseReactionRate(reverse_rate)`, are to be called.
  - Function input variables:
    – `bulkChemistry (AbstractChemistry*)`
    – `cellChemistry (AbstractChemistry*)`
    – `currentBulkConc (const std::vector<double>&)`
    – `currentCellConc (const std::vector<double>&)`
3. `GetReactionType()`
  - Function description: This function returns a string type which provides the name of the reaction type; for example returning `MassActionCoupledMembraneReaction`. This function may also be used for reaction tracking purposes, but it is currently used in the `MembraneTablet` function to set the name of the membrane rate type. This name needs to be the name in which the reaction files label the reaction type in order for the correct class calls to be made.
4. `UpdateReaction()`
  - Function description: This is a void function with no inputs provided for the case that a reaction’s behaviour needs to alter outside of the `React()` function. Possible utilities include a switching of behaviour in transport style based on concentrations, time, or system properties.
5. `ParseReactionInformation()`
  - Function description: This function parses the parameters and variables needed to process the transport process. In terms of the written file reaction, these data values occur in the space after the ; delimiter. This string is to be parsed into the data values by the user using a string delimiter. The user needs to identify the delimiter of this string as a member value in this membrane reaction class. The values parsed are to be also stored as member values and may be utilised in the `UpdateReactionRate()` function where necessary.
  - Input variables:
    – `reaction_information (string)`
    – `IsReversible (bool)`

### S6 Derivation of Schnakenberg parameter sets

The Schnakenberg reaction system involves chemical species U and V that are produced, inter-converted, and removed via the reactions

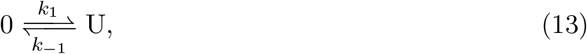

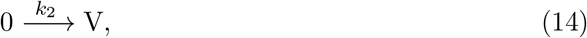

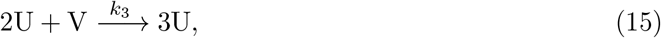

where *k*_1_, *k*_−1_, *k*_2_, *k*_3_ denote reaction rate constants. Applying mass action kinetics to these reactions yields the ODE system

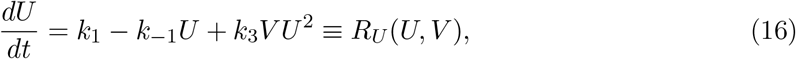

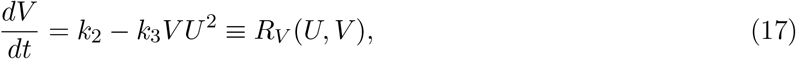

for the concentrations of U and V. The Schnakenberg reaction-diffusion system extends this model to include diffusion of U and V with constant diffusion coefficients *D_U_* and *D_V_*, respectively, leading to the set of coupled PDEs

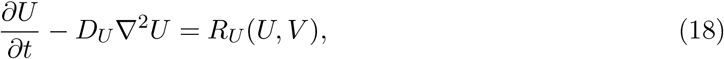

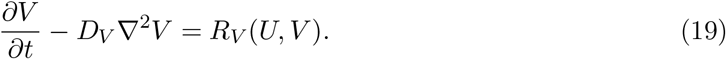

The Schnakenberg model is a spatial case of the Gierer-Meinhardt activator-inhibitor model (Gierer and Meinhardt, 1972). Following previous mathematical analyses (Murray, 2003; Korvasová *et al*., 2015; Guin *et al*., 2012; Gambino *et al*., 2013), we present necessary conditions for pattern formation via diffusion-driven instability (DDI) in this model. For a DDI to occur, we require the spatially uniform solution to (18)–(19) to be linearly stable in the absence of diffusion, but unstable in the presence of diffusion. In the absence of diffusion (*D_U_* = *D_V_* = 0), equations (18)–(19) have the unique steady-state solution

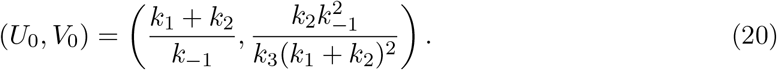

By requiring both eigenvalues of the Jacobian to have negative real part, we find that (20) is linearly stable in the absence of diffusion if

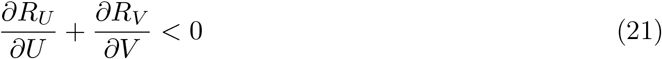

and

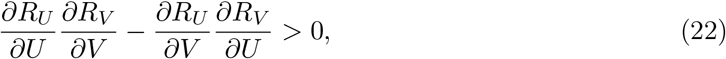

where the partial derivatives of *R_U_* and *R_V_* are evaluated at (20). By requiring at least one eigenvalue to have negative real part, we find that (20) becomes linearly unstable in the presence of diffusion if

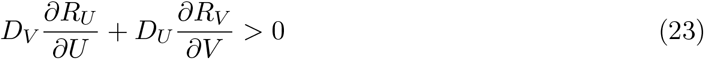

and

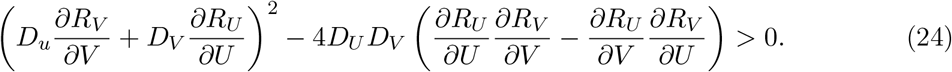

After some algebra, we find that the above conditions correspond to the following inequalities on the reaction rate constants *k*_1_, *k*_1_, *k*_2_, *k*_3_ and diffusion coefficients *D_U_*, *D_V_*:

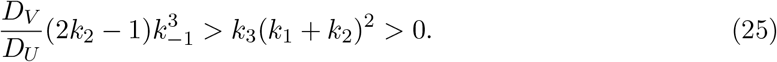

